# Sentences, Words, Attention: A “Transforming” Aphorism for the Discovery of pre-miRNA Regions across Plant Genomes

**DOI:** 10.1101/2022.07.14.500029

**Authors:** Sagar Gupta, Vishal Saini, Rajiv Kumar, Ravi Shankar

**Affiliations:** Studio of Computational Biology & Bioinformatics, The Himalayan Centre for High-throughput Computational Biology, (HiCHiCoB, A BIC supported by DBT, India), CSIR-Institute of Himalayan Bioresource Technology (CSIR-IHBT), Palampur (HP), 176061, India; Biotechnology Division, CSIR-Institute of Himalayan Bioresource Technology (CSIR-IHBT), Palampur (HP), 176061, India; Academy of Scientific and Innovative Research (AcSIR), Ghaziabad, Uttar Pradesh-201002

**Keywords:** microRNA, Transformers, CNN, Deep learning, Genomics

## Abstract

Discovering pre-miRNAs is the core of miRNA discovery. Using traditional sequence/structural features many tools have been published to discover miRNAs. However, in practical applications like genomic annotations, their actual performance has been far away from acceptable. This becomes more grave in plants where unlike animals pre-miRNAs are much more complex and difficult to identify. This is reflected by the huge gap between the available software for miRNA discovery and species specific miRNAs information for animals and plants. Here, we present miWords, an attention based genomic language processing transformer and context scoring deep-learning approach, with an optional sRNA-seq guided CNN module to accurately identify pre-miRNA regions in plant genomes. During a comprehensive bench-marking the transformer part of miWords alone significantly outperformed the compared published tools with consistent performance while breaching accuracy of 98% across a large number of experimentally validated data. Performance of miWords was also evaluated across *Arabidopsis* genome where also miWords, even without using its sRNA-seq reads module, outperformed those software which essentially require sRNA-seq reads to identify miRNAs. miWords was run across the Tea genome, reporting 803 pre-miRNA regions, all validated by sRNA-seq reads from multiple samples, and 10 randomly selected cases re-validated by qRT-PCR.

## Introduction

miRNAs are prime regulatory small RNAs (sRNAs) having ∼21 bases length which post-transcriptionally regulate most of the genes and stand critical for most of the processes of eukaryotic cells including their development and specialization (1). Mature miRNAs are derived from longer precursor miRNA molecules (pre-miRNAs) which are double-stranded RNAs (dsRNAs) with terminal hairpin loop. Discovering these pre-miRNAs is the central to the problem of finding miRNAs and genomic annotations for them. However, finding the pre-miRNAs remains a challenge, and more so in plants. Unlike animals, in plants mature miRNA formation from the precursors is a single step process. Also in terms of complexity, secondary structures of plant pre-miRNAs are way more complex and longer than those of animal pre-miRNAs (2), making them difficult to detect accurately. The traditionally considered sequence and structural properties and features to identify miRNAs are also responsible for difficulty in identifying them as they display lots of variability across the plant genomes in terms of sequence composition, structural, and thermodynamic properties.

A comparison of these traditional properties properties like AU%, GC%, length of sequence, number of bulges, terminal loop length, minimum free energy (MFE), maximum bulge size, mismatches and stem length for pre-miRNAs in plants and animals display a good amount of variability and overlap with other genomic elements for several of these properties miRNAs and non-miRNAs regions for the same properties (Figure 1 and Supplementary Table S1 Sheet 1). This all suggest that how they are prone to wrongly identify the miRNA regions.

**Figure 1:**
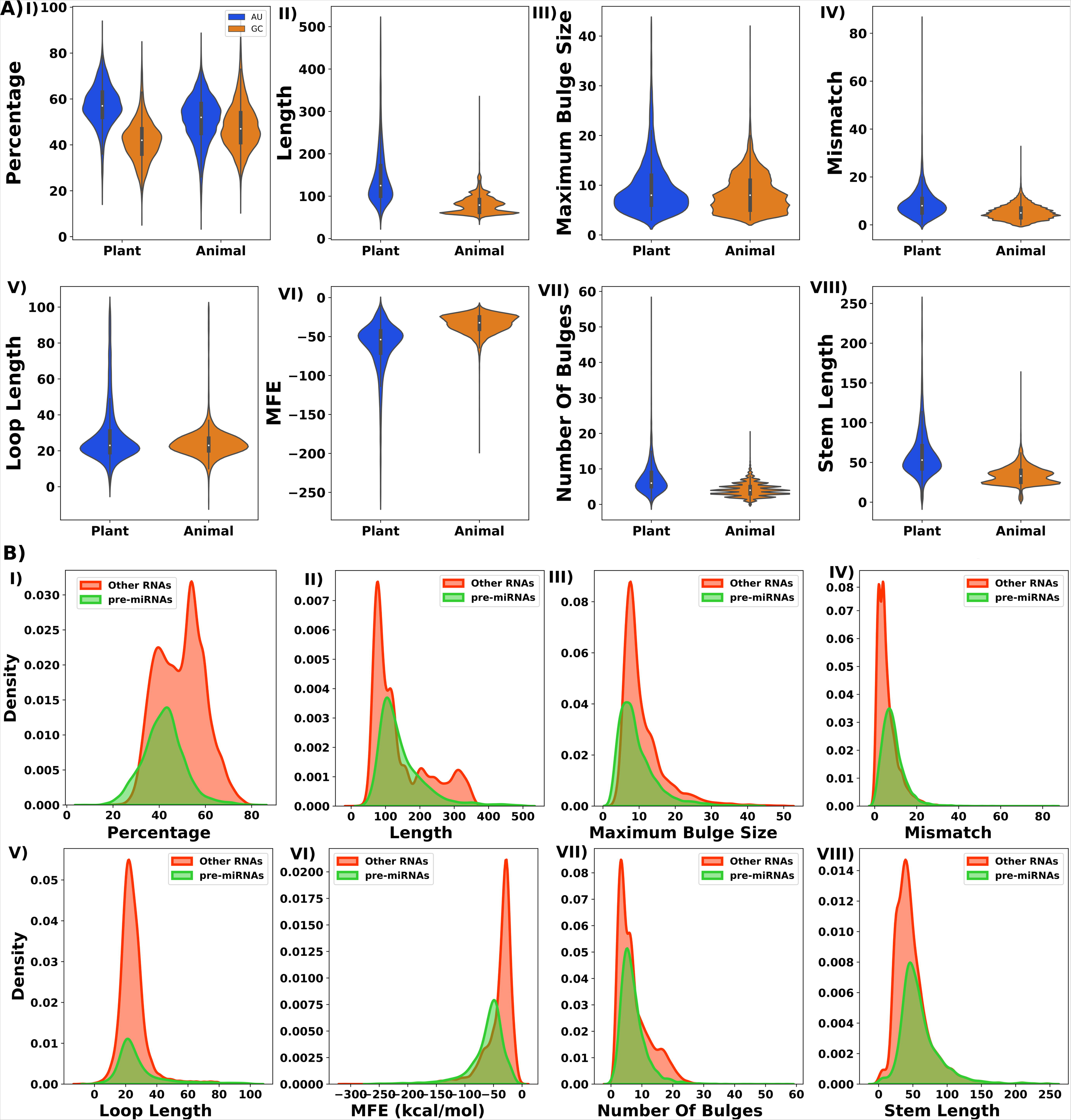
Distribution pattern of traditionally considered properties for miRNA characterization. **A)** Pattern of distribution comparison between animals and plants pre-miRNAs. Values differ a lot between animals and plants as unlike animal pre-miRNAs, plants pre-miRNAs display much more complexity and variability. **B)** Pattern of distribution comparison between pre-miRNAs v/s other RNAs in plants. As can be seen clearly that most of these properties are actually not strong discriminators as lots of overlap in their values occur between pre-miRNAs and other RNAs.

The difficulties in the identification of plant miRNAs has been so much that it could be fathomed from the fact that it has been plagued of huge volume of false reportings, and in year 2018 miRBase had to scrap a large number of the reported plant miRNAs data (3). An urgent call was made to critically look into the process of plant miRNA identification and annotation process, following which in the some important suggestions were made (4). These studies highlighted the need of support by sRNA-seq for reporting novel miRNAs and seriously questioned the capabilities of existing software pool to identify plant pre/miRNAs which were flooded with false predictions and false positive reporting. The same studies, therefore, also recommended that credibility of any such software must be judged by using it across some well established and annotated genomes like *Arabidopsis* and evaluate their false positive rates.

Though experimental techniques like direct cloning and quantitative real-time PCR (qRT-PCR) are used identify miRNAs with high expression levels, they are costly and cumbersome for genomic level identifications and remain mainly limited for experimental validation purpose or studying a handful of miRNAs. Off-late, experiments like sRNA-seq and arrays studies have been identified as the main way to identify and profile miRNAs/pre-miRNAs across the genome, but they too essentially require support from computational approaches while using the sequencing read data mainly as support guide while in actual at their core they use some pre-miRNA discovery algorithms. These approaches have their own shortcoming and they are not immune to false identifications. First, their dependence on sRNA-seq data makes them not approachable by all and a costlier stuff as one will have to first manage the sequencing experiments and costs related to that. Secondly, these all methods also utilize the same old features and rules which we described above while describing how they work as an inefficient discriminators. Third, they can capture only those pre-miRNAs which express in any given condition, and miRNA expression is highly specific. Many of their rules are not suitable and may even capture non-miRNAs as well as discard genuine miRNAs.

These pre-miRNA discovery algorithms have evolved with age. In the identification of miRNA precursors, identification of secondary structure patterns, hairpin loops, and their thermodynamic stability have been the most followed approaches. Additionally, homology and conservation patterns were also used to locate similar kind of precursors in other genomes. Compared to the conservation and rules based methods, machine-learning-based methods are mainly anchored on sequence and structure based features of pre-miRNAs with more mature automated statistical learning process, sans the rules based approaches. They have specially emphasized upon the RNA secondary structure and hairpin loops identification. Although, groups like Bentwich *et al* suggested that there are about 11 million hairpins in human genome, making it a daunting task to correctly identify miRNA precursor candidates from hairpins (5). Shallow machine learning approaches like support vector machines, tree based, Bayesian, and ensemble learnings have grossly marked the process of machine learning based approaches for pre-miRNA discovery. However, majority of these developed methods were specifically designed to identify animal pre-miRNAs but rarely for plant pre-miRNAs. Only tools like Triplet-SVM (6) were tested for plant pre-miRNAs. As the plant pre-miRNAs differ greatly from the animal pre-miRNAs and are more complicated, plant miRNA discovery kept lagging while lots of software were developed for animals. Yet, plant miRNA biology took forward the Machine-learning revolution caused by Triplet-SVM with tools like PlantMiRNAPred (SVM based) (2), HuntMi (Random Forest based) (7), MiPlantPreMat (SVM based) (8), and plantMirP (Random forest) (9). Table 1 highlights the various categories of core algorithms employed to discover pre-miRNA regions across genome. As apparent grossly, the area of plant miRNA discovery rarely witnessed any major entry of deep-learning based plant pre-miRNA discovery tools until very recently.

**Table 1:** List of software for pre-miRNAs identification.

| S. No. | Software | Algorithm | Year | sRNA-Seq | Webserver (W)/Standalone (S) |
| --- | --- | --- | --- | --- | --- |
| 1 | MiRFinder | SVM | 2004<br>(10) | N | S |
| 2 | MIRcheck | Target<br>Identification | 2004<br>(11) | N | S |
| 3 | FindMiRNA | Kmer based<br>sequence<br>similarity | 2005<br>(12) | N | S |
| 4 | MiMatcher | SVM | 2005<br>(13) | N | S |
| 5 | PalGrade | Scoring<br>hairpins by<br>thermodynamic<br>stability and<br>structural<br>features | 2005<br>(5) | N | - |
| 6 | Triplet-SVM | SVM | 2005<br>(6) | N | S |
| 7 | MicroHARVESTOR | Sequence<br>similarity | 2006<br>(14) | N | S |
| 8 | RNAmicro | SVM | 2006<br>(15) | N | S |
| 9 | miPred | Random forest | 2007<br>(16) | N | W/S |
| 10 | microPred | SMOTE | 2009<br>(17) | N | W/S |
| 11 | miRanalyzer | sRNA-Seq<br>based filtering | 2009<br>(18) | Y | W/S |
| 12 | MIReNA | sRNA-Seq based filtering | 2010 (19) | Y | S |
| 13 | mirDeep-P | sRNA-Seq based filtering | 2011 (20) | Y | S |
| 14 | PlantMiRNAPred | SVM | 2011 (2) | N | S |
| 15 | miR-BAG | Bagging ensemble (SVM, BF-Tree and Naive Bayes) | 2012 (21) | Y | W/S |
| 16 | mirDeep2 | sRNA-Seq based filtering | 2012 (22) | Y | S |
| 17 | mirDeep* | sRNA-Seq based filtering | 2013 (23) | Y | W/S |
| 18 | HuntMi | Random forest | 2013 (7) | N | S |
| 19 | ShortStack | sRNA-Seq based filtering | 2013 (24) | Y | S |
| 20 | MiPlantPreMat | SVM | 2014 (8) | N | S |
| 21 | miR-PREFeR | sRNA-Seq based filtering | 2014 (25) | Y | S |
| 22 | plantMirP | Random forest | 2016 (9) | N | S |
| 23 | DP-miRNA | Boltzman machines based deep learning | 2017 (26) | N | S |
| 24 | deepSOM | DL based self organizing maps | 2017 (27) | N | W/S |
| 25 | deepMiRGene | LSTM | 2017 (28) | N | S |
| 26 | miRNAss | Semi-supervised and transductive learning | 2018 (29) | N | S |
| 27 | DeepMiR | CNN | 2019 (30) | N | S |
| 28 | mirDNN | Convolutional deep residual networks | 2021 (31) | N | S |

Very recently, deep learning (DL) techniques have been implemented successfully to tasks such as speech and image recognition, largely eliminating the manual construction of features and their engineering. They have been very effectively digging out better but hidden features for model building which are otherwise difficult to detect manually (32). DL approaches like convolutional neural networks (CNN) and recurrent neural networks (RNN), the two dominant types of DNN architectures which have shown huge success in image recognition and natural language processing (NLP) (33, 33, 34, 35). CNN takes input in the form of pixeled matrices where it effectively compresses the features to detect the spatial patterns. Natural language processing approaches like recurrent neural nets (RNN) were developed for processing sequential data (35). RNNs evolved further into long short-term memory (LSTM) approach where short distanced associations within a sequence could be detected and memorized. miRNA biology has very recently witnessed some of them for pre-miRNA discovery, which though not developed exclusively for plants, but may be trained on the plant specific data also.

However, despite the forays into DNN based software to detect pre-miRNAs, there remains lots of voids to be filled. First is the inconsistent performance where huge gaps were found when benchmarked across different datasets. Secondly, most of them are still based on direct reading of the input sequences and work with 4-states nucleic acid sequence inputs and 3 states secondary structure inputs. Third, barring CNN, DL approaches like LSTM and RBM deep learning approaches require lots of compute power, time, and resources. They can’t be parallelized and become very high on compute resources requirements and time to train the network. Further to this, they fail to detect the associations effectively if the sequence length is increased or long distanced associations are present. As already mentioned above, the plant pre-miRNAs differ significantly from animal miRNAs from sequence to structural properties, as can be seen from the Figure 1. Plant pre-miRNAs have much larger sequences than animals, which may complicate learning. And above all, the existing software pool at present still fail miserably in their practical application of genome annotation for miRNAs, as noted by some recent studies (4). Recently, a new revolutionary DL architecture, Transformers, has been introduced in the area of artificial intelligence, which has emerged as a highly efficient architecture for language processing tasks (36). It uses self attention mechanism on the input which can be processed in parallel while more effectively capturing the long distanced associations and contexts within any sequential data.

Inspired by these milestones in Deep Learning, the present work proposes a novel deep learning system where the transformer defines the first phase whose sequence output from its encoder becomes the input to a shallow learning approach, XGBoost, which makes the classification decision score. The important aspect is that unlike most of the existing deep-learning approaches, miWords sees a genome sequence as a set of sentences composed of words made from monomers, di-mers, and pentamers capturing sequence, structure, and shape based information and communication among themselves. Besides this, genome is also seen as a sentence pool made of words from structural triplets from the RNA secondary structures. Context wise their association including long ranged ones are successfully detected which is otherwise missed by the existing software pool for miRNA discovery. A comprehensive benchmarking study was performed whose results showed that miWords outperformed all the compared software with highly significant margin, just based on this transformer part alone.

The real application of such software in genomic sequence annotation of pre-miRNA discovery has been a challenging part where most of the existing software for pre-miRNA discovery generate lots of wrong classification besides being lethargically slow to scan genomic sequences. The existing software pool hardly considers the long distanced relative standing of the genomic regions to characterize pre-miRNA regions, which is one big reason why they end up producing lots of false positives. We noticed that the regions having miRNAs generate a specific transformer scoring (T-score) pattern when compared to the non-miRNA regions, and this could work as a strong discriminator which could further boost pragmatic miRNA discovery with genomic context information. The T-score generated by the transformer across the genomic region forms a scoring plot which captures the relative standing of the scores for various regions on which a CNN module was trained. This CNN part can scan the regions and their T-scores in a relative and contextual manner for the genome wide T-scoring profile. It remarkably lowers the false identifications which is otherwise hardly seen with any existing approaches. In addition to this CNN module, one more optional CNN module has been provided where the user can supply the sRNA-seq data to further enhance the accuracy. Thus, a user can run miWords with and without sRNA-seq support data, and in both the ways can get highly accurate results.

Considering the 2018 guidelines for miRNA discovery and annotations which asks to prove the performance of such software across well annotated genome like *Arabidopsis*, to measure the degree of false positives in the real application, we also ran miWords across the whole genome of *Arabidopsis* and carried out the genomic annotation performance measure while comparing with other existing tools. miWords outperformed all of them with just 10 false positives. Even without considering sRNA-seq read data, miWords outperformed all those well established software which essentially require sRNA sequencing read data, making it much affordable besides being a better performer. Finally, miWords was applied across the Tea genome whose miRNAs annotation is still not well established. It identified a total of 803 pre-miRNA regions across the Tea genome, all of which were supported by sRNA-seq reads and 10 randomly selected candidates also re-validated with RT-qPCR for their transcriptional status. miWords has been made freely available as a source code as well as web-server. We expect miWords to drastically improve the scenario of plant genomes annotations and plant miRNA biology.

## Materials and Methods

### Datasets Sources

In this study data was retrieved for 27 different plant species. For the 27 plants species, the positive set included experimentally validated pre-miRNAs and negative datasets contained mRNAs, rRNAs, snoRNAs, snRNAs, tRNAs, and other non-coding sequences while following the same protocol which we had followed in animal pre-miRNA discovery tool miRBAG (21). The pre-miRNAs sequences and their co-ordinates were fetched from miRBase version 22 for all the available plant species (37). Species which had no genome information were discarded, bringing the total number of species to 27. From Ensembl Plants (v51) and NCBI negative instance sequences were downloaded for same selected 27 plant species. The repeated sequences were filtered out. After that, the same number of sequences were extracted randomly for each species with respect to their species pre-miRNAs number. A combined positive and negative datasets were created for all these 27 plants species to build a model that could act as a universal classifier for plants pre-miRNAs.

### Dataset Generation

Uniform length sequences with flanking regions were obtained for every individual positive instance which varied pre-miRNA-wise. For creating the positive dataset, the central base of the terminal loop was treated as the reference point, a standard protocol (21). This placed the reference point appropriately at constant position, providing uniformity across all possible dataset instances, irrespective of variation in length and loop size. Considering the central base of the terminal loop as the midpoint, genomic sequences up to 200 bp were extracted from the flanking regions.

For the negative dataset creation, the terminal loops were identified in the hairpin structures of the negative instances. As followed above, the central base of the terminal loop was mapped as the reference central position for the creation of every negative instance, as in case of the precursors for reference position identification. The negative dataset consisted of different types of RNA sequences including ribosomal RNA, small nucleolar RNA, small nuclear RNA, transfer RNA, mRNA and long non-coding sequences, all taking pseudo-hairpin shapes. For all the instances, the genomic co-ordinate of central reference position was considered for taking 200 bp sequence with equal flanks. With this approach, a consistent sequence length for positive and negative instances was maintained. These sequences were used to create the training and testing datasets for all 27 species. In the entire study this dataset is known as Dataset “A”.

Some observations have been made in past that a substantial fraction of miRBase entries may not be a genuine miRNA. Considering such data may alter the quality of the dataset (3). Therefore, we performed a filtering step on the above mentioned dataset “A” to get high confidence positive instances. We retrieved pre-miRNAs data from sRNAanno (143 species) (38), PmiREN (87 species) (39), and PNRD (150 species) (40). sRNAanno uses majorly three tiered stringent filtering criteria based on sequencing reads data, sequence/structural rules and strands expression ratio to filter highly confident miRNA candidates. Very similar to sRNAanno, PmiREN also employs sequencing reads data support to annotate a pre-miRNA while also considering their functional potential to target genes using PARE-seq data. miRNAs annotated in PNRD are also high confidence and filtered one. PNRD uses recurring expression profiles across various tissues, experimental evidences and reportings, literature support, target support besides the sequence and structure based characterization to annotate a miRNA candidate. Thus, all the above mentioned pre-miRNA information sources provide high confidence miRNA annotations which could be used to vet the miRBase data to zero upon the high confidence miRNA candidates. Data retrieved from these three databases and miRBase (v22) were merged together and a cut-off of 95% similarity was applied to remove similar sequences with CD-HIT-EST (41) in the dataset. Later, only those pre-miRNAs form miRBase were considered which had support from at least one of these databases. Those sequence’s without single support were discarded. In this way, a refined dataset was created. In the entire study this dataset has been called as Dataset “B”.

We additionally constructed an independent dataset, known as Dataset “C”, for another layer of unbiased benchamarking. This dataset is based mainly on the instances considered by other tools considered for their benchmarking purpose while covering 75 plant species. Like Dataset “A”, the positive instances were from the pre-miRNAs available at miRBase. But unlike Dataset “A” which has positive instances from only 27 species whose coordinates are available as discussed earlier, Dataset “C” had also those pre-miRNAs from miRBase for the species whose genomic coordinates/information are presently not available. Negative instances of this dataset were also built from the dataset for the same for the selected tools. Redundant sequences were dropped. For the negative instances, a total of 92,000 instances were collected for this dataset. Dataset “C” was purely used for objective comparative bench-marking on which all compared tools were trained and tested on common training and testing datasets. Besides these datasets, one more version of each of the datasets “A” and “B” were created where all sequences from *Arabidopsis thaliana* were removed. This was done exclusively for the benchmarking study across the *Arabidopsis* genome in order to avoid any chance of memory and bias.

In addition to this all, another reference dataset for *Arabidopsis* was retrieved from the work of Bugnon, et al (2021) where they had benchmarked various tools for miRNA identification recently (42). They had focused on the performance of the tools on the real situation data like genomes where class imbalance is pronounced with much higher instances of non-miRNA instances. This dataset had induced class imbalance with much larger number of negative instances (13,56,616) than the positive one (839), collected from *Arabidopsis thaliana* genome.

### Encoding of sequence data

The Encoder-Decoder architecture with Transformer has emerged as one of the most effective approaches for the neural machine translation, sequence-to-sequence, and binary classification. The prime importance of the method is the potential to train a single end-to-end model directly on source and target sentences having the capability to handle variable length input and output sequences. However, deep networks like transformers, LSTM, and RNN work by performing computation on integers, passing in a group of words won’t work. So these input sequences were tokenized for further computation. Provided a character or word sequence and a defined vocabulary set, tokenization is the procedure of cutting it up into unique numerical units, called tokens. These tokenizers processes words from the sentence as input and output a unique numerical representation for the tokenized word which becomes input for the embedding layer of the model. Tokenization followed by embedding layer allowed to vectorize the words into a fixed sized (28 elements each) vector of numeric values. The process of tokenization was implemented using TensorFlow (Keras) Tokenizer class for end-to-end tokenization of the positive and negative datasets. We created the Tokenizer object, providing the maximum number of words as our vocabulary size, which we had in the training data. Tokenizing the data while mapping the words to unique numeric representation, the vocabulary and words within the genomic sequences were encoded.

To encode sequences, every single instance was transliterated into its various words in terms of monomeric, dinucleotide, and pentameric sequences in an overlapping window manner (43). Dinucleotides and pentamers provide structural stacking and shape information, respectively (44). The secondary structural information of these RNA sequences were also used. This information of structure for each sequence was obtained by RNAfold of ViennaRNA Package v2.4.18 (45). RNAfold determines the secondary structure of RNA and gives an output in dot-bracket form (“(“,”.“,”)”). To tokenize the information obtained from RNAfold, the dot-bracket symbols were transliterated in the following manner: (“(“−>”M”, “.”−>”O”, “N” −>”)”. For a sequence length of 200 bases, a maximum length of the input vector was of 793 elements. The encoded monomers, dinucleotides, pentameric sequences and secondary structural words were independent of each another. In this way, the network would determine the associations between the sequence derived inputs and the structure triplets based inputs in its own way. The encoded instances were then fed as input the transformer encoder to train and evaluate the models.

### Implementation of the Transformers -XGBoost classification system

With the tokenized sequences, encoded vectors were used to build models to classify and distinguish pre-miRNAs from other genomic elements using a deep-shallow learning approach: The multi-headed attention system of Transformer’s encoder which derives the most confident contextual associations and places them into a hidden space vectors, which in turn becomes the input for the XGBoost part to classify and generate the classification score (T-score). Both were implemented using python scikit-learn, XGBoost, Keras, and Tensorflow libraries. In addition, as per the standard practice, the dataset was broken into 70% and 30% as train and test sets, and using the 70% training part the model was built and tested upon the 30% totally untouched unseen testing part. Further, in order to assess the consistency of the approach and its observed performance, 10-fold randomized trials were performed where 10 times the entire data-set was randomly split into 70:30 training and testing data, and new model was built and tested from the scratch every time. Also, this was ensured that absolutely no instance overlapped between the train and test set in any fold. This ensured fair training and testing process without any scope of memorization of instances.

The input layer consists of embedding and position encoding layers which operate on matrices representing a batch of sequence samples. Embedding encodes each word ID (unique token number) into a word vector whose length is the embedding size, resulting in a (samples, sequence length, embedding size) shaped output matrix. For any given genomic sequence of length “*l*” (sentence size), there are “*n*” words in it. Embedding of these arranged words can be represented as:

Let the sentence S = {w_1_, w_2_,……, w_n_}

where, w**_n_** = n^th^ word in the sentences. Every such word in a sentence is converted into a vector of *d-dimension* whose elements (I_d_) carry the optimized numeric weights:

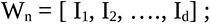

Therefore, the sentence can be represented in the form of embedded words matrix X:

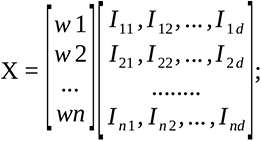

where, each row corresponds to the word in “S”. This matrix has, thus a dimension of *n x d* (number of words in the sentence x the dimension used to represent each word in embedding vector).

Each word of the matrix “X” is also combined with its corresponding positional embedding “P”. The position embedding has same dimension “d” as is for the word embedding vector W_n_:

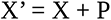

where, P = [p_1_, p_2_,….., p_d_] and the values of P are derived using the following equations:

P_d_ = *sine*(*index*|10000*^index^*/^*no. of dimensions*^)for all the even positions in the vector;.

P_d_ = cos (*index*|10000*^index^* ^/*no*^*^. of dimensions^*)for all the odd positions in the vector.

Position encoding produces a similarly shaped matrix that can be added to the embedding matrix. Shape of the matrix (samples, sequence length, embedding size) produced by the embedding and position encoding layers is maintained throughout the Transformer, which is finally reshaped by the final output layers. The input embedding layer sends its outputs into the next layer. Similarly, the output embedding layer feeds into the next layer (encoder layer).

The encoder passes input into a multi-head attention layer. The attention module consists of one or more attention heads. The attention module splits its query, key, and value parameters N-ways and feds each split independently through a different head and then merged together to generate a final attention score. This entire process of attention score generation has five major steps:

**Step 1:** From the above mentioned input matrix X’ derived from the embedded words create the Query matrix (Q), Key matrix (K), and Value matrix (V):

Q = X’.W_O_, where W_Q_ is the optimizable weight matrix for the query matrix generation.

K = X’.W_K_, where W_K_ is the optimizable weight matrix for the key matrix generation.

V = X’.W_V_, where W_V_ is the optimizable weight matrix for the value matrix generation.

All these three weight matrices are randomly initialized.

**Step 2:** Inner product between Query (Q) and Key (K) transpose matrices: Q.K^T^

This step establishes the weights for association between the words within a sentence and captures their dependence.

**Step 3:** Scale the Query-Key inner product to stabilize the gradients:

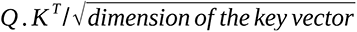

**Step 4:** Normalization through softmax function, which ensures that values are in the range of 0-1:

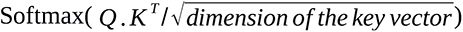

**Step 5:** Compute the attention matrix “A” to get the attention score for each word in the sentence: This is achieved by taking inner product of the above mentioned softmax normalized query and key inner product with the Value matrix (V):

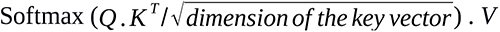

This is a single column vector which holds the attention score for each positional word and their relative closeness in the given sentence.

For multi-headed attention, the same steps were repeated according to the number of heads and their individual attention scores vectors were finally concatenated and forwarded to the block of feed-forward network of the transformer encoder for further processing. Figure 2 provides a snapshot of how this entire system is working. The output from multi-head attention layer passed into dropout layer which helps to reduce over-fitting, which is followed by a layer of normalization. Afterwards, the output from this layer passes to a feed-forward layer, which then sends its output to the next dropout layer in the stack. A feed-forward layer was implemented gathering its input from the previous layer, followed by a layer of GlobalAveragePooling1D which then passes its output to the third Dropout layer of the model. The Dropout layer passes output into the fully connected hidden layers.

**Figure 2:**
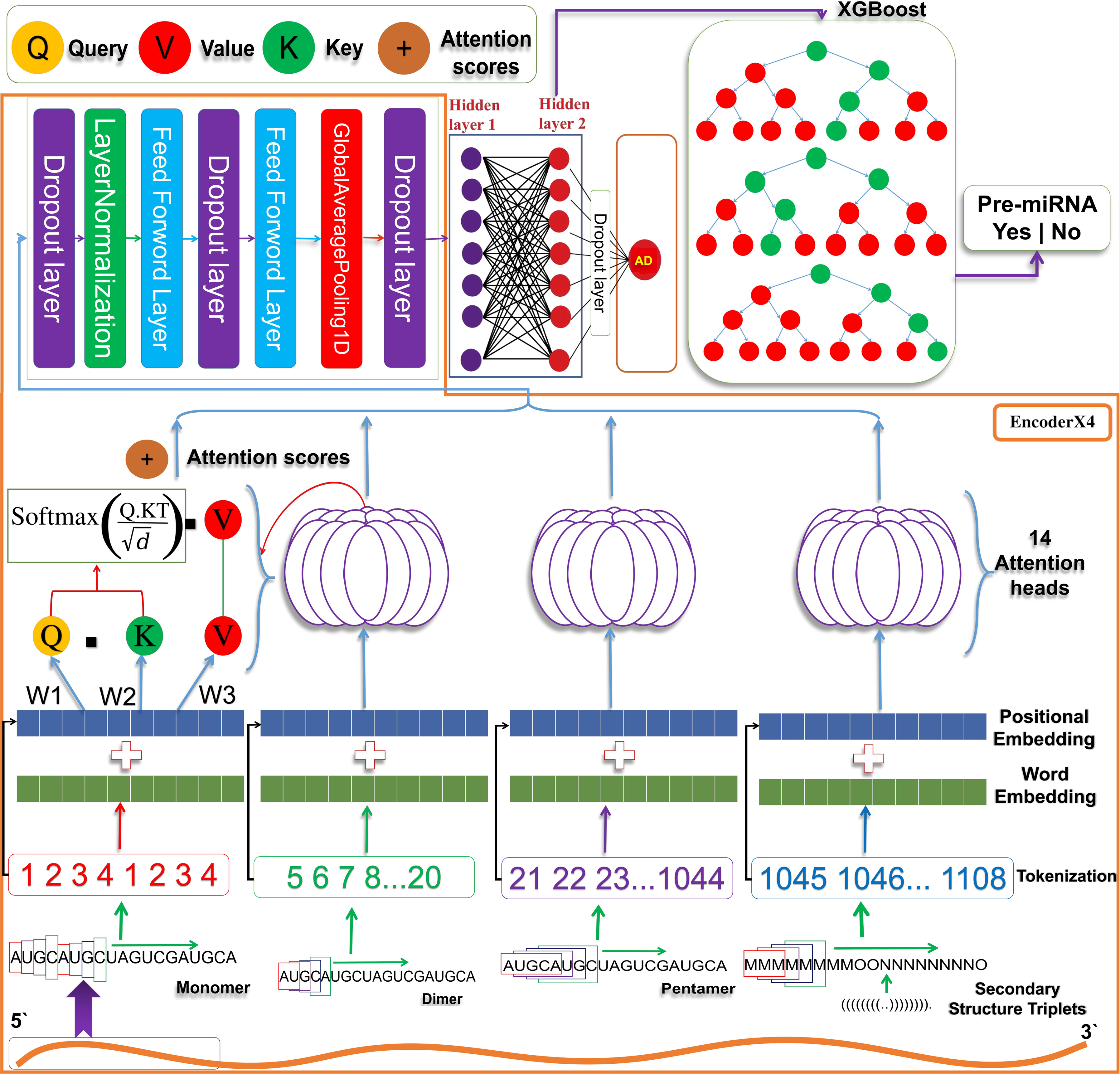
Implementation of the transformer based module to identify pre-miRNAs. The image provides the brief outline of the entire computation protocol implemented to develop the Transformer-XGBoost based model to identify pre-miRNAs. This illustrates how a genomic sequence can be seen as a sentence composed of words and their related arrangements which can be efficiently learned through multi-headed transformers. The various nucleotides k-mers and RNA secondary structure triplets define the words for any given regions (the sentence). The words and their attention scores are evaluated through query, key, and value matrices which are then passed to different layers of deep-learning protocol to present its learning for classification job through XGBoost.

The performance of the Transformer was evaluated for a numbers of hidden layers where finally total two hidden layers were found performing the best and the connections between the nodes were made dense. For the model, the number of nodes across the two hidden layers were tuned. All the component layers were optimized for their suitable numbers and component nodes number by iterative additions. Different activation functions were examined for the layers from a pool of available activation function. The fourth Dropout layer takes from the last hidden layer and passes its output into an LeakyReLu activation function based single node output classification layer. Binary cross entropy loss function was used to calculate the loss. “Adadelta” optimizer was used at this point to adjust the weights and learning rates. Adadelta adapts learning rates based on a moving window of gradient updates instead of accumulating all past gradients even when many updates have been done. The learning rate was set to 0.583 for the optimizer and the model build was trained using 20 epochs and batch size of 40 instances. The transformer part derived the hidden features and their relationships which got structured also, on which classification could be done in much superior manner. Since the present problem in this study was not translation but classification, decoders were not needed and instead the encoder output were taken as input for next step of extreme gradient boosting. For this purpose the output of the transformer was passed to the XGBoost classification part. The number of encoders and performance were studied and it was found that no significant major change in performance was observed by increasing the encoder layers, and it only dropped after 6^th^ encoder layers. Thus, we continued with single encoder layer to keep it lighter while giving the similar performance. However, for Dataset “B” based learning four encoder layers were found best performing and model based on Dataset “B” were implemented with four encoders, while all the rest hyper-parameters remained the same.

### Optimizaiton of Tansformer-XGBoost system

A gradient boosting framework, XGBoost, is a decision-tree-based ensemble Machine Learning algorithm which has been consistently rated at the top in shallow learning approaches at Kaggle bench-markings. In the classification phase with XGBoost, grid search was applied for parameter optimization using scikit-learn function RandomizedSearchCV. Following hyper-parameters were optimized with the grid search: “eta/learning rate”, “max_depth”, “objective”, “silent”, “base_score”, “gamma”, “subsample”, “eta”, “colsample_bytree”, “n_estimators”, “min_child_weight”, eval_metric”, “tree_method”, “reg_alpha”, “reg_lambda”. Gradient boosted decision trees learn very quickly and may overfit. To overcome this, shrinkage was used which slows down the learning rate of gradient boosting models. Size of the decision tree were run on different combinations of max-depth. Values changed until stability was gained as the logloss got stabilized and did not change thereafter. The final max_depth value was 6.

The final model obtained was saved in hierarchical data format 5 (HDF5). Since the entire system is implemented here using TensorFlow and scikit-learn, the HDF5 format provided the graph definition and weights of the model to the TensorFlow structure and saved the model for classification purpose. Each and every hyper-parameter values involved to finalize this hybrid model were fixed using an in-house developed script which tested various combinations of values of the hyper-parameters to pick the best ones. This entire optimization process was done using two different approaches: Random search optimization and Bayesian optimizations. Figure 2 shows the detailed workflow of the implemented architecture.

### Performance Evaluation

The performance of the built model was evaluated. Four classes of the confusion matrix namely true positives (TP), false negatives (FN), false positives (FP), and true negatives (TN) were evaluated. The performance of the raised transformer based model was assessed the performance metrics like sensitivity, specificity, accuracy, F1-Score, and Mathew correlation coefficient (MCC). Sensitivity/True Positive Rate (TPR) defines the proportion of positives which were correctly identified as positives. The specificity value informs about the proportion of negative instances correctly identified. Precision defines the proportion of positives with respect to total true and false positives. F1-score measures the balance between precision and recall. Besides these metrics, Mathew’s Correlation Coefficient (MCC) was also considered. MCC is considered among the best metrics to understand the performance where the score is equally influenced by all the four confusion matrix classes (true positives, false negatives, true negatives, and false positives) (46). A good MCC score is an indicator of robust and balanced model with high degree of performance consistency. AUC/ROC and mean absolute error were also measured for the build model.

Besides this all, the consistency of performance on the developed approach was evaluated through 10-fold random independent trials of training and testing. Every time the dataset was randomly split into 70:30 ratio with first used to train and second part used to test, respectively. Every single time data was shuffled and random data was selected for building new model from the scratch. Accuracy and other performance measure were calculated for each such model. In order to avoid any sort of imbalance, memory, and bias, it was ensured that no overlap of instances existed ever between the train and test sets.

Performance measures were done using the following equations:

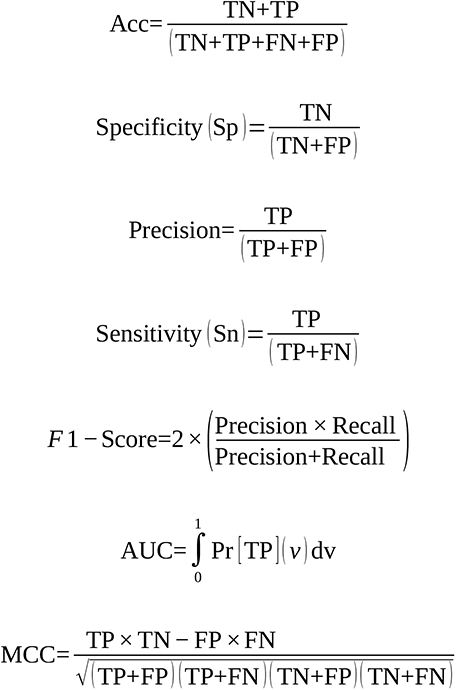

Where:

TP = True Positives, TN = True Negatives, FP = False Positives, FN = False Negatives, Acc = Accuracy, AUC = Area Under Curve.

### CNNs based implementation of genomic scanning capabilities using Transformer-XGBoost scoring profiles

Two different CNN modules were constructed for the identification of pre-miRNAs most confident potential regions across the genomes. First one is genome wide T-scoring based CNN model and the second one is the optional one which uses Read Per Million (RPM) based CNN to further improve the identification of pre-miRNA regions across the genomes while using sRNA-seq reads data as the guide.

The first module works with the Transformer modules scoring output for every genomic position. Thus, the scanned genomic sequence is transformed into its corresponding transformer scores sequence for each scanned position until the last window frame. It appears like a graph plot where the T-scoring around the pre-miRNA regions appear different than the regions not having them. This T-scores sequence is converted into a one hot encoding acting as an input to a convolution layer. The scoring profile input has a dimension of 280×10. If the length of some instance was found shorter it was padded for the empty columns with value of zero. Size of 280 covers the base positions in a window, for each of which corresponding T-score exists. 10 dimensions come from 10 different ranges of T-scoring ranging from 0 to 1. To evaluate the performance of the scoring based CNNs, various number of hidden, convolutional, maxpooling, and batch normalization layers were tested and finally two convolutional, one maxpooling, four batch normalization, and four hidden layers were applied in a fully connected manner. The number of the nodes across both the dense hidden layers were tuned based on the number of filters used in the convolution layer. All the component layers were optimized for their best numbers and component nodes number by iterative additions. Additionally, the kernel size and strides were optimized by trying different values in an incremental order. A sigmoid activation function based single node classification layer was used with binary cross entropy loss function to calculate the loss. “Adam” optimizer was used at this point to adjust the weights and learning rates. The batch size was set to 4 and the number of epochs was set to 25.

The second CNN based module is optional as it requires availability of short reads sequence mapping information. It takes RPM value for each base in the input while the input genomic sequence is represented in the form of RPM value sequence. For this RPM based second module RPM profiles were created and transformed later into a vector of length of 280 elements, where each element holds the scaled normalized sRNA read depth value for the position of nucleotide in the sequence window. Here also, padding was done for the shorter input instances. Similar to the T-scoring based CNN, here also various numbers of hidden, convolutional, and maxpooling layers were tested and finally one convolutional, one maxpooling, two batch normalization, and two fully connected hidden layers were found best performing. The number of the nodes across both the dense hidden layers, the number of filters used in the convolution layer, the kernel size and strides were optimized by trying different values. Similar to the above mentioned T-Score CNN, a sigmoid activation function based single node classification layer was used with binary cross entropy loss function to calculate the loss. “Adam” optimizer was used at this point to adjust the weights and learning rates. The batch size was set to 64 and the number of epochs was set to 25.

To train and test the T-scoring based CNN module, two different dataset were created from Dataset “A” and “B”, respectively. The first dataset was created from the known pre-miRNA regions of *Oryza sativa* taken in Dataset “A” and the second dataset was created from the pre-miRNA regions of *Oryza sativa, Glycine max, Arabidopsis thaliana,* and *Zea mays* after refinement (Dataset “B”). For both datasets, the 500 bases from their 5’ and 3’ ends were extracted from the genome, which acted as the negative instances which could help recognize the boundaries and shift towards the corresponding miRNA region. All these sequences were represented as corresponding T-score for each base position. Same was done for the pre-miRNA regions also. As discuused earlier, the sequence which is transformed into its corresponding probability scores appears like a graph plot where the scorings around the pre-miRNA regions appear different than the flanking regions not having them. For the first dataset (Derived from Dataset “A”), the entire data from *Oryza* was considered, where the total number of positive instances (pre-miRNA regions) was 604, and the total number of non-pre-miRNA regions (the 500 bases flanking sequences around the pre-miRNAs) was 604. Similarily, for the second dataset, entire refined data (Dataset “B”) for *Oryza sativa, Glycine max, Arabidopsis thaliana,* and *Zea mays* was considered, where the total number of positive instances (pre-miRNA regions) was 631 positive instances, and the total number of non-pre-miRNA regions (the 500 bases flanking sequences around the pre-miRNAs) was 631 negative instances. Later, scores were converted into one hot encoding representation, acting as the input to a convolution layer with a dimension of 280 × 10.

Similarily, with the RPM based CNN module the extracted sequences from the genome of *Oryza sativa, Glycine max, Arabidopsis thaliana,* and *Zea mays* were mapped back to the genome to calculate its reads per million (RPM) value for every single base. A total of 201 different samples covering a total of 66 experimental conditions and a total of seven billion sRNA-seq reads (239-GB) were considered for raising the RPM CNN module. The fully annotated *Arabidopsis thaliana* genome version (GCA 000001735.1 TAIR10) was used as the test set to benchmark the performance for genomic annotation. It has to be noted that for the benchmarking on *Arabidopsis thaliana* genome and associated datasets, sequences from *Arabidopsis thaliana* were removed from the models of both T-scoring based CNN module and RPM based CNN module. For these Arabidopsis specific T-scoring and RPM based CNN models two different models were raised exclusively from Dataset “A” (model “A”) and “B” (model “B”) where instances from Arabidopsis were completely removed to avoid any chance of bias and memorization.

### Optimization of CNNs

In the T-scoring based CNNs model, the T-scores are converted into a one hot encoding as the input to a convolution layer and has the dimension of 280×10, where the various scoring bins defined the first dimension and the base positions in the length of 280 window defined the second dimension. The input layer was followed by a convolution layer containing 64 channels at each position with 2×2 kernel size. The input sequence was padded if the length was shorter than 280 in order to ensure a constant size of the input matrix. The output resulted into a dimension of 279 × 9 × 64 representation after the convolution. This passes into a MaxPooling layer. This layer included 32 nodes having the kernel size of 2×2. Max-pooling helped in reducing the dimensions of convoluted sequence into a dimension of 139×4×64 which is then flattened by the flatten layer. The output from flatten layer passes into first dense layer which is followed by Batch Normalization. This layer was used to overcome the over-fitting problem during training process. Likewise, the output from previous layer passes into second dense layer and then into second Batch Normalization layer. Similarly, the input passes through two more combinations of dense and Batch Normalization. It goes finally into the output layer with 50 dimension. The output layers had a node with sigmoid activation function. The model was compiled by binary cross entropy loss function to calculate the loss which was optimized with “Adam” optimizer, for a batch size of four and 25 epochs. The model produced probability score for every instance passed.

In the optional RPM based CNNs model, the scaled normalized RPM values of each base became the input to a 1D convolution layer with dimension of 280×1. The convolution layer had 32 channels at each position with kernel dimension of of 5×1. The output from the convolution layer passed into a Max-pooling layer whose output was flattened and converted into a flatten layer. The output from the flattened layer passed into two fully connected dense layers. Later, it was followed by a final output layer with a sigmoid activation function. The model was compiled by binary cross entropy loss function to calculate the loss which was optimized by using “Adam” optimizer with a batch size 64 and epochs of 25. The model produces probability score for every instance passed.

Another model was raised combining T-scoring based CNNs and RPM based CNNs into a bi-modal CNN. This model system unlike the above one, processes T-score sequence and RPM profile sequence in parallel and combined manner, and has been raised when the sRNA-seq read data is available and to be used essentially. To construct this bi-modal CNN, previously built architecture for T-scoring and RPM based CNN were used. Both the architectures were joined together with a concatenate layer after they passed through their respective last batch normalization layer before connecting to the output layer. The output from the concatenation layer passes to the final output layer with a sigmoid activation function. The model was compiled by mean absolute error loss function to calculate the loss which was optimized by using “Adam” optimizer with batch size 4 and epochs of 250. The model produces probability score for every instance passed. To train and test this bi-modal CNN module, the same section of dataset “B” was used from T-score and RPM based CNN models on which they were trained and tested previously (covering seqeunces from *Oryza sativa, Glycine max, Arabidopsis thaliana,* and *Zea mays* after refinement (Dataset “B”)).

### Performance benchmarking for genomic annotations and application demonstration

To identify the pre-miRNAs on *Arabdiopsis thaliana* and *Camellia sinesis*, we downloaded both the genomes from NCBI (‘GCA 000001735.1 TAIR10’ and ‘GCA 004153795.2 AHAU CSS 2’) for performance benchmarking for genome wide pre-miRNAs annotation and as the application demonstration, respectively. The genomes were scanned by the transformer module through an overlapping sliding window of 200 bases up to n-200^th^ position. The generated position-wise scores sequence was scanned through an overlapping sliding window of 280 elements, where every window becomes the input to the CNN modules as described above.

*Arabidopsis* genome annotation for miRNAs were obtained from miRBase (v22). Seven different published tool’s annotations for *Arabidopsis* (miRanalyzer (18), MIReNA (19), mirDeep-P (20), mirDeep2 (22), mirDeep* (23), ShortStack (24), miR-PREFeR (25)) were also considered for the corresponding comparative performance measure process (3, 4).

### Validation of the identified pre-miRNAs candidates using sRNA-seq reads

For the validation of the identified pre-miRNAs, the sRNA reads were considered by mapping them to the genome. These sRNA-seq fastq data were collected from GEO and SRA databases which had 88 and 104 different read files for *Arabdiopsis thaliana* and *Camellia sinesis,* respectively (Supplementary Table S2 Sheet 1-2). Genomic sequences, annotations and reference RNA sequences were downloaded from NCBI. Trimmomatic v0.39 (47) and in house developed reads processing tool, filteR (48), were used to filter out poor quality reads, read trimming, and for adapter removal. Filtered reads were mapped back to the genome using Hisat2 (49). Complete list of various conditions and sources is available in Supplementary Table S2 Sheet 1-2 . To remove any bias and noise due to some random elements, two different criteria were applied: (i) Reads which appeared more than five times in any given experiment were only considered, and (ii) only those mapping regions were considered which got support from at least for two different experimental conditions. All these reads were subjected to validation for identified pre-miRNAs across the genomes. The co-ordinates of the short reads were obtained from the mapped results and were intersected it with results obtained from miWords utilizing bedtools (50). Since many miRNAs are also homologous, the identified miRNAs by miWords were searched for homology support using BLAST with already reported plant miRNAs in miRBase. Related information about sRNA reads file is provided in Supplementary Table S2 Sheet 1-2.

### Server and Stand-alone Implementation

The entire server was implemented in Apache-Linux platform using HTML5 and PHP. Majority of the codes were developed in python and shell for data curation and back-end processing. Statistical processing and calculations were implemented were also executed using modules developed with python. The standalone version was developed in python and shell. The entire work was carried out in the Open Source OS environment of Ubuntu Linux platforms.

### RNA isolation and quantitative real-time analysis

Total RNA was isolated from tea leaves (Camellia sinensis) from the CSIR-Institute of Himalayan Bioresource technology (32° 05’ 59’’N; 76° 34’ 04’’ E; 1305 m a. s. l) experimental farm (CSIR-IHBT-269) (51). Approximately 10-12 plants more than eight years old were randomly selected from three sub-populations from the same farm, thus representing three biological replicates for leaf sampling. Samples were harvested in liquid nitrogen and stored at -80 °C for RNA isolation. Total RNA was isolated from leaf tissue (100mg) using Trizol (Invitrogen, USA) according to (52). Total RNA was treated with RNase-free DNase I (Invitrogen, USA) as per as manufacturer’s protocol. The cDNA synthesis was performed using random hexamer and SuperSrcipt® III Reverse Transcriptase (Invitrogen, USA), as per as manufacturer’s protocol. Primers of 10 pre-miRNAs were designed using primer 3 software v.0.4.0; Applied Biosystem (Table 2). Quantitative real time-PCR was performed using the standard protocol on Applied Biosystem, USA. In brief, 2.5 µg of the 1/100 dilution of cDNA with water was added to 5.5µl of SYBR green (Thermo scientific, USA), 2.5nM each primer, and water to 10 µl reaction mixture. Amplification was performed with an initial denaturing at 95 °C for 7 min, followed by 40 cycles of 95 °C for 10s, 53 °C for 30 s, and 72 °C for the 30s. Relative expression of each pre-miRNA was calculated using the equation 2^-ΔC^_T_ where ΔC_T_ = (C_TPre-miRNA_ – C_T18SrRNA_) (53). 18S rRNA was taken as an internal control to normalize the variance in cDNA. To simplify the relative presentation expression of each pre-miRNA was multiplied by 10^6^. All reactions of qRT-PCR were performed using three biological and three technical replicates.

**Table 2:**
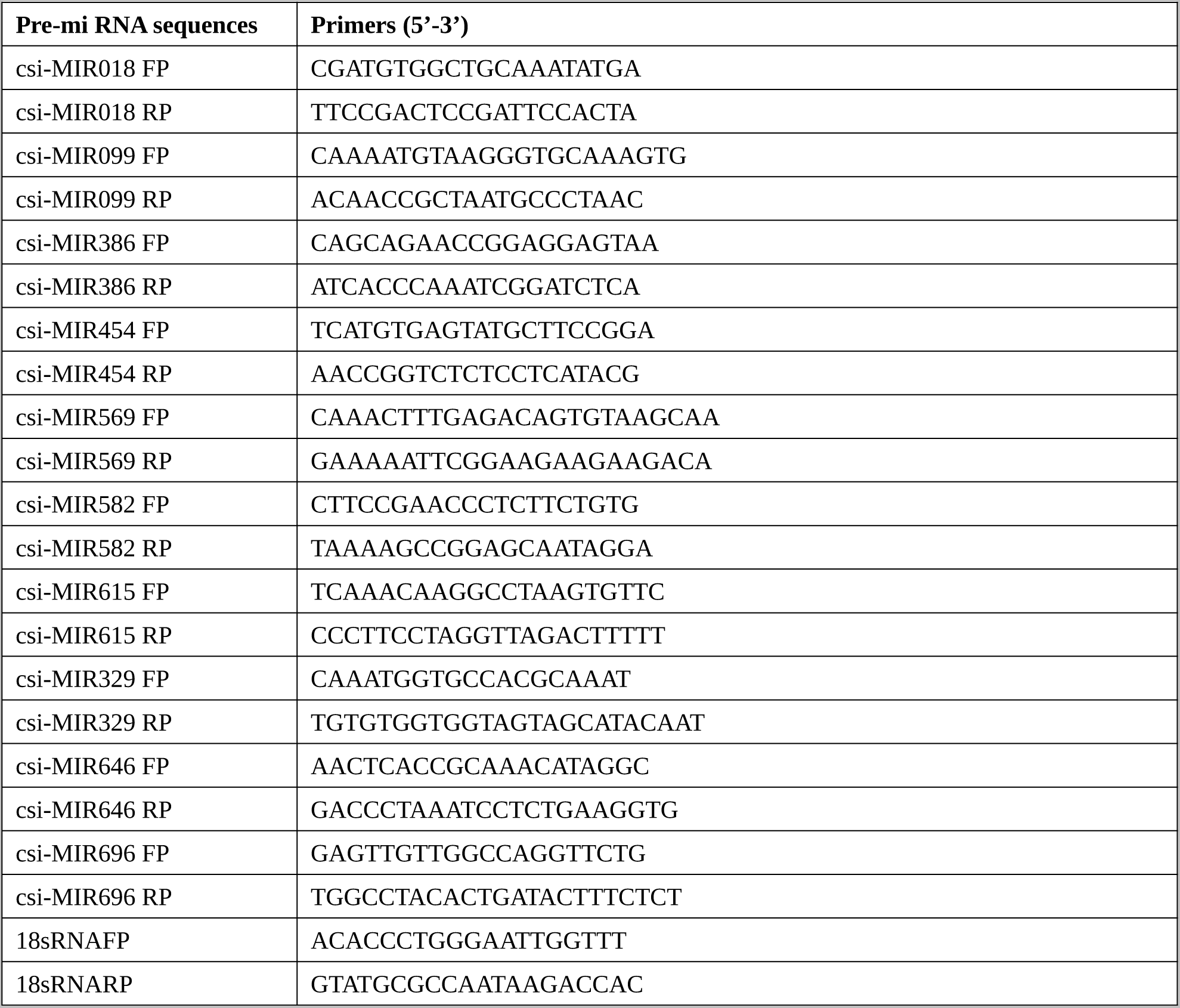
Primer list for 10 selected Pre-miRNAs from Tea genome taken for revalidation through qPCR.

## Results and Discussion

### The datasets

Presently, there are 8,615 known plant pre-miRNAs in the miRNA database miRBase v22 (http://www.mirbase.org/). After the retrieval of data, only those species sequences were kept which had their respective genome information and those having no information were discarded from the pool. 5,685 pre-miRNAs belonged to 27 plant species which formed the initial positive dataset. Supplementary Table S2 Sheet 3 shows the pre-miRNAs distribution across the 27 plant species. To construct the negative dataset, same number of instances were collected from the species selected for their corresponding positive dataset. For only 14 out of 27 species, various non-coding, coding RNAs were available at Ensembl plants (v51) from which a total of 5,684 RNAs of different classes were retrieved while eliminating the redundant sequences. This dataset has been called Dataset “A” in the present study.

The dataset “B” was constructed from miRBase (v22) (37), sRNAanno (38), PmiREN (39), and PNRD (40) for construction of a dataset with high confidence pre-miRNA instances as many entries in miRBase have been under argument as not proven correctly as miRNAs. In material and methods section above details about sRNAanno, pmiREN, and PNRD has already been given and how they ensured high confidence positive instances in the present study. By eliminating sequences of >95% similarity with CD-HIT-EST (41) and considering sequences from miRBase only when at least any one of these three databases supported a pre-miRNA instance, a total of 3,923 pre-miRNAs qualified as high confidence positive instances of pre-miRNAs. The equal amount of negative instances were taken randomly from the Dataset “A”, and together this dataset formed the high confidence Dataset “B”.

Besides the above mentioned datasets, another Dataset “C” was built exclusively for testing and objective comparative bench-marking purpose. This dataset became the source for common training and testing datasets to attain the objective comperative benchmarking where all the compared tools have been trained and tested on this common dataset. This dataset was built from the datasets used by the different tools like HuntMi (7), miPlantPreMat (8), PlantMiRNAPred (2), and plantMirP (9). A total of 16,404 pre-miRNAs sequences were retreived, covering 75 plant species. After removing similar sequence, 9,214 plant pre-miRNAs (positive instances) remained. Similarily, 92,000 RNAs of different classes (negative instances) were retrieved after removal of the redundant sequences. Figure 3 provides the illustrated details on all these datasets.

**Figure 3:**
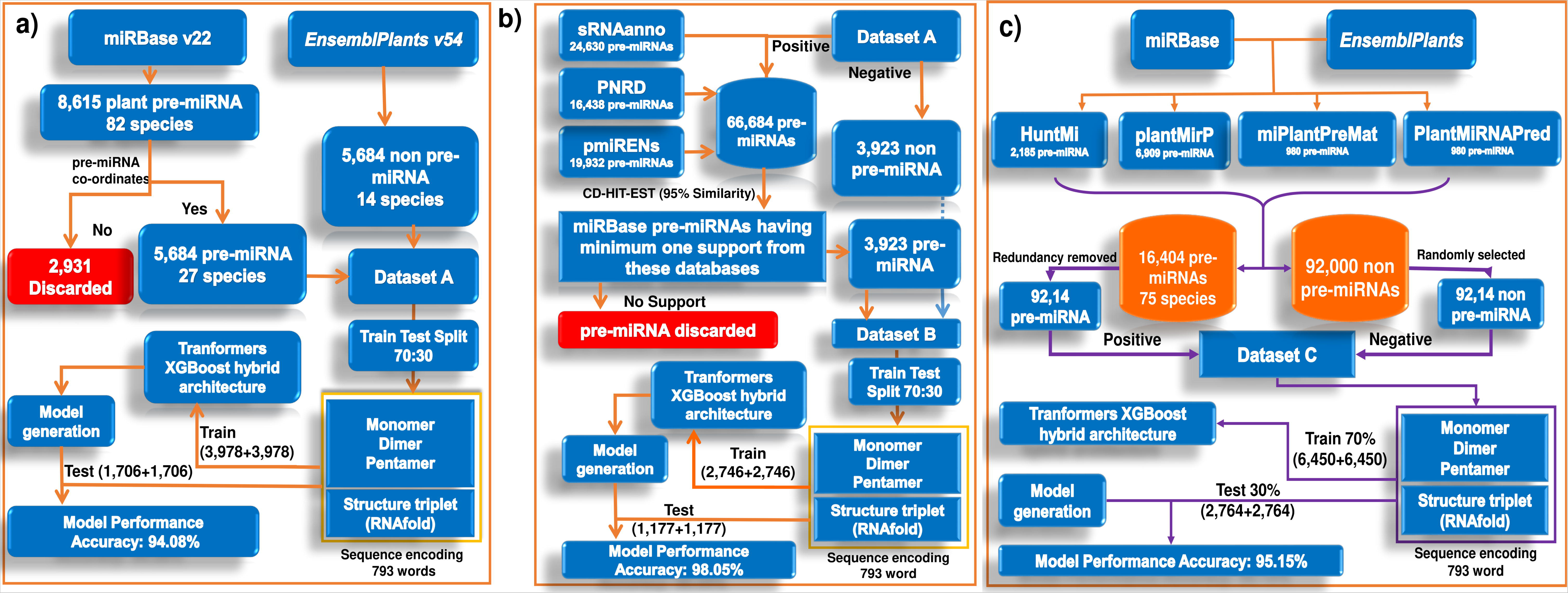
Flowchart representation of dataset processing and formation. a) The protocol followed for Dataset “A” creation, b) Protocol followed for Dataset “B” creation, and c) Protocol followed for Dataset “C” creation.

In addition to this, to fathom the performance on the real situation data like genomes where class imbalance is pronounced with much higher instances of non-miRNA instances, dataset provided in the study by Bugnon et al (2021) was also considered (42).

### Sentences, Words, and Attention! Seeing genome as a pool of sentences through transformers delivers high accuracy

Most of the existing pre-miRNA discovery tools depend upon some traditionally identified feature sets highly focused on the hair-pin loop structures and sequence composition. They build around properties like Minimum free energy (MFE), stem length, AU/GC content, pairing in stem, terminal loop size etc, most of which are inherited from miPred (17). However, these properties exhibit significant differences between animal and plant system, and within plants themselves, these properties exhibit lots of variations. Figure 1 shows the distribution plots for some of such features to build the pre-miRNA models which exhibit a lot of difference from animals as well as variation among themselves and overlap with other types of RNAs. Also, in most of the existing tools, there is absolutely no effort made to record their relative standing and context which largely limit their practical application when used to annotate genomic sequences (4, 54).

One of the above cited studies clearly showed how poorly most for the existing software to detect pre-miRNAs perform when they face real situation application of performing genomic annotation. They recommended that compared to the traditional machine-learning approaches, it is the need of the time to focus upon the development of the methods based on DL approaches which may perform better than the other machine learning methods. Considering these seminal works and limitations of the existing machine learning approaches, the current study proposes a revolutionary transformers deep-learning based approach where context and relative standing of the properties have been emphasized upon to come up with a highly accurate and practical pre-miRNA discovery system, miWords.

For the building of a universal model for plant pre-miRNA regions for its characterization against the other types of RNAs, we used 13 different combinations for various sequence input encodings: 1) Monomers, 2) Dimers, 3) Trimers, 4) Pentamers, 5) Structure triplets, 6) Monomers+Dimers, 7) Monomers+Trimers, 8) Monomers+Dimers+Trimers, 9) Monomers+Dimers+Pentamers, 10) Monomers+Dimers+Pentamers+Triplets, 11) Monomers+Dimers+Trimers+Triplets, and 12) Monomers+Dimers+Trimers+Pentamers+Triplets. They were evaluated for performance through the raised transformer encoder based model. An assessment was made for each encoding considered where the Dataset “A” was split into 70:30 ratio to form the train and test dataset. The model was trained with the train set and all the above mentioned properties encodings were done accordingly. This protocol came in action as an ablation analysis to evaluate how each of these individual encodings of the sequence was contributing towards the building of model for accurate classification. The sequences were taken in a uniform length of 200 bases while considering the reference from the midpoint of the terminal loop, as described in the methods section.

At the first, the Transformer was trained and tested without the XGBoost gradient boosting to evaluate its performance. First, the test of performance was done with monomeric sequence representation and its corresponding encodings fed into the input layer of the Transformer. The observed accuracy for monomeric encodings was just 72.36%. This was followed by feeding of dimeric, trimeric, and pentameric sequence representations and their encoded sequences into the input layer of the Transformer. This returned an accuracy of 73.21%, 75.36%, and 79.01%, respectively, while covering a total of 199, 198, and 196 words, respectively.

The reasoning for considering monomeric representation was that they capture sequence composition. While the dinucleotide densities representation has been proven very useful to reflect the base stacking and secondary structural tendencies (45, 55). Pentamaric sequences are reflective of the nucleic acids shape which determine protein-nucleic acids interactions (44, 56). All these are critical to characterize a pre-miRNAs. Besides the above mentioned sequence based properties, the secondary structure stem-loop based features were also used for the representation and encoding, as pre-miRNAs exist in the stem-loop hairpin form. RNAfold derived stable secondary structure of the considered region was used for the structural representation. For the sequence encoding the extracted dot-bracket secondary structure of these sequences were converted as following: (“(“−>”M”, “.”−>”O”, “N” −>”)”. These were fed into the input layer in the form of triplet words covering a total of 198 words per sentence. This fetched an accuracy of 77.09%. As can be seen here, individually all these properties did not score much and needed information sharing with each other. To obtain encoding of a particular type, the sequence was broken down into sub-sequences in an overlapping fashion to gather all the existing possible combinations which were later converted into encodings.

Above evaluation results showed varying influences on pre-miRNA identification observed for the different encodings. The next step was observing the influence of combining these sequence and structure derived words encodings and learning on them. Combining of the encodings was done in a gradual manner in order to see their additive effect on the classification performance. These combination of encodings yielded a better result than using any single encoding. Combining monomers with dinucleotides (399 words) yielded an accuracy of 81.23% while the combination of monomers + trimers (398 words) yielded an accuracy of 84.06%. In addition, the combination of monomers + dimers + trimers (597 words) and monmers + dimers + pentamers (595 words) achieved the accuracy of 87.63% and 89.32%, respectively. As we know, secondary structure holds critical role in miRNA biogenesis, combining these encodings with structure triplets based encodings led to further superior result. Monomers+dinucleotides+trinucleotides+structure triplets (795 words), Monomers+dinucleotides+trinucleotides+pentanucleotides+structure triplets (991 words), and monomers+dinucleotides+pentamers+structure triplets (793 words) combinations yielded the accuracy of 93.67%, 93.84%, and 93.96%, respectively, with the latter one having the better balance between the sensitivity and specificity values. Thus, combinations of different representations and encoding for the genomic sequence markedly improved the performance through the natural language processing approach of transformers. Figure 4 presents the plots for the accuracy, sensitivity and specificity values distribution observed for the various combinations of the sequence encodings.

**Figure 4:**
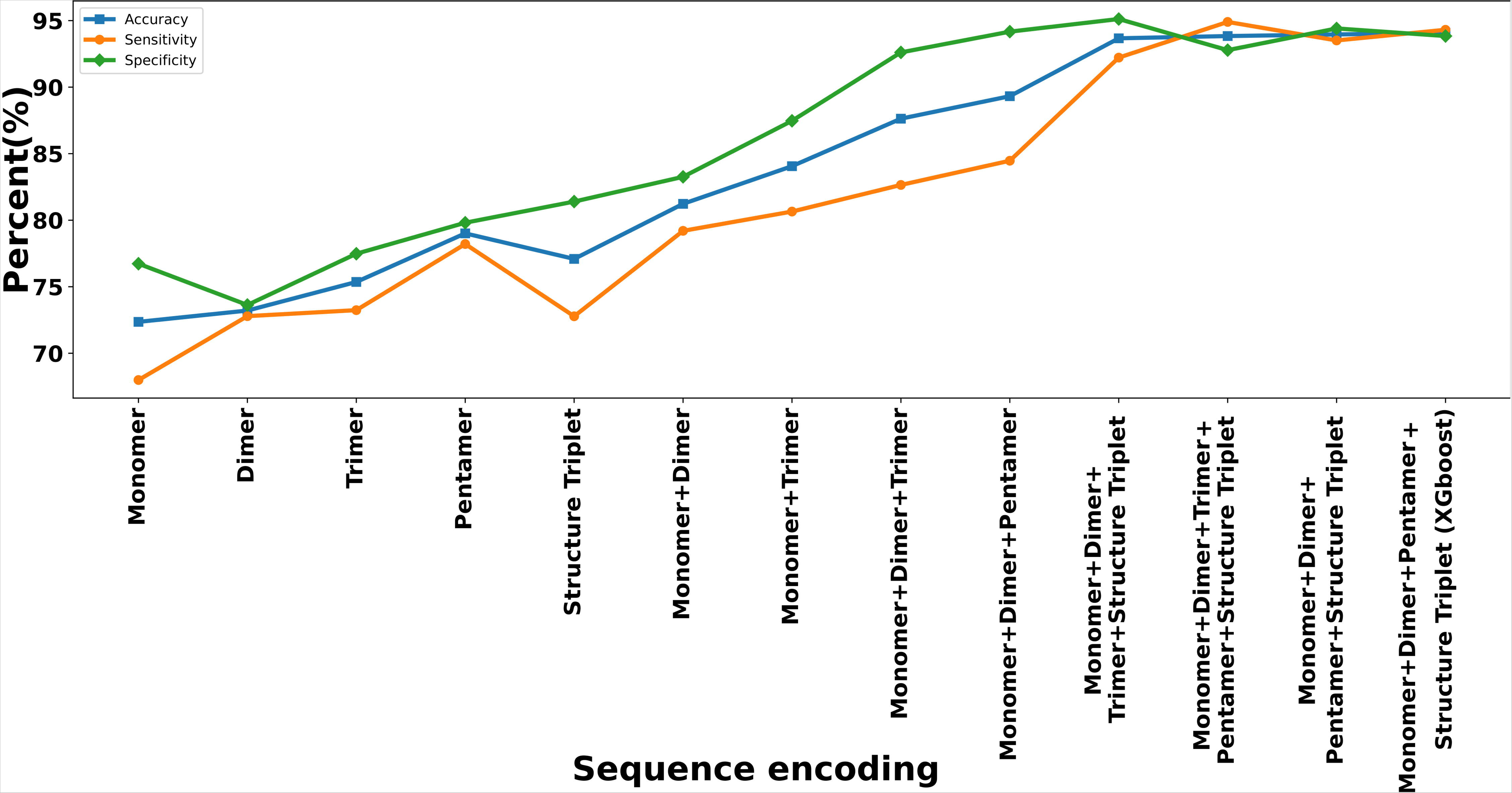
Ablation analysis for five main properties in discriminating between the negative and positive instances. Impact of combination of the monomer, dimer, trimer, pentamers, and structure triplet properties based sequence encodings. These encodings appeared highly additive and complementary to each other as the performance in accurately identifying pre-miRNAs increased substantially as they combined together.

The Transformers built from the combination of monomer+dimer+pentamer+structure triplet encodings delivered a good accuracy of 93.96%. There was a gap of 0.9% between sensitivity and specificity, though not a big gap, yet we tried to reduce it further. In doing so, the output layer of the transformer having the LeakyReLu activation function was replaced by XGBoost for the classification purpose. XGBoost was the choice as it has come consistently at the top along with deep learning approaches in Kaggle benchmarkings, and performs exceptionally good on structured data and manually extracted features input sets. In our case, the Transformer’s encoder became the feature feeder to XGBoost. This also strengthened the performance further while leveraging from the two best and different approaches of machine learning. This hybrid deep-shallow model reduced the performance gap between the sensitivity and specificity to just 0.46% while also increased the accuracy slightly to 94.08% (Supplementary Table S3 Sheet 1). Likewise, another model was derived form Dataset “B” which was based on high confidence positive instances. This model attained an accuracy of 98.04% on its test set along with specificity of 98.56% and sensitivity of 97.54%. This all became the first part of the transformer based pre-miRNA identification system, which can even work independently and can be used directly for pre-miRNA regions identification. It was further enhanced for pragmatic genomic scanning and annotation purpose which is discussed in the upcoming sections.

### Optimization of the Transformer-XGBoost system

Optimization of the hyper-parameters is an important step to derive the best possible model. The transformer part had encoder role which learned across the hidden space of features and presented it to the classification part done by XGBoost part. The transformer encoders had multi-headed attention layer where a total of 14 self attention heads were found best performing. Multihead attentions help a transformer to avoid misunderstanding the relationships between the words by multiple vetting by the different transformer heads for the derived attention scores. The input sequence was padded if the length was shorter than 200 bases in order to ensure a constant size of the input matrix. This output was passed into a dropout layer with dropout fraction 0.1. By passing - dropout fraction, 10% of the hidden units were randomly dropped during the training process of the model. This layer helped to reduce over-fitting. Later, this output was normalized by the second layer called normalization layer. This layer was followed by a third layer called Feed Forward layer with 14 nodes and followed by second Dropout layer with dropout fraction of 0.1 and second Feed Forward layer with 14 nodes. The output from Feed forward layer passed into GlobalAveragePooling1D layer, followed by the third dropout layer with dropout fraction of 0.16. Next to this layer, pooled feature maps were passed to two fully connected layer. The hidden layers in the present study had two dense layers with both having 38 and 12 hidden nodes with RELU and SELU activation function, respectively. The hidden layer output was passed into the fourth dropout layer in the stack with dropout fraction of 0.17 which passed its output into the last layer. Finally, the output of the dense layer with 12 dimension was passed to the last and final output layer, a node with LeakyReLu activation function. The model was compiled by binary cross entropy loss function to calculate the loss which was optimized by using the “Adadelta” optimizer with learning rate of 0.583. An accuracy of 94.08% and 98.05% was observed for the test sets from Dataset “A” and “B”, respectively.

In the classification part of the hybrid Transformer-XGBoost, XGBoost takes input from the second fully connected layer of the Transformers stack. Grid search was applied for hyperparameter optimization using scikit-learn function RandomizedSearchCV. Following hyperparameters were finalized after the grid search: params = {”eta/learning rate”: 0.22, “max_depth”: 6, “objective”: “binary:logistic”, “silent”: 1, “base_score”: np.mean(yt), “gamma”: 6.4, “subsample”: 0.6, “eta”: 0.4, “colsample_bytree”: 0.83, “n_estimators”: 1400, “min_child_weight”: 4.76, “eval_metric”: “logloss”, “reg_alpha”: 149.468151996443, “reg_lambda”: 0.02399001301159498, “tree_method”: ’approx’}. To overcome over-fitting shrinkage was used which slows down the learning rate of gradient boosting models. At the value of 6, stability was gained as the logloss got stabilized and did not change thereafter. The output from the XGBoost returned the probability score (T-Score) for each input sequence. The probability score indicated the confidence of each instance as non-pre-miRNA or pre-miRNA. If the T-Score >0.50, the corresponding input sequence was identified as pre-miRNA else a non-pre-miRNA.

The final hyperparameters set for the output layer of the implemented model was: {“Activation function”: LeakyReLu, “Loss function”: binary crossentropy, “Optimizer”: Adadelta}. The related information about optimization towards the final model is listed in Supplementary Table S3 Sheet 2-3 and illustrated in Supplementary Figure S1.

### Consistent performance across different validated datasets reinforces miWords as a universal classifier for plant pre-miRNAs

As mentioned in the Methods section, for performance testing three different datasets, “A”, “B”, and “C” were created. Dataset “A” had 5,684 positive instances and 5,684 negative instances, totalling 11,368 instances. Dataset “B” had 3,923 positive instances and 3,923 negative instances, totalling 7,846 instances. 70% of dataset A and B were used for training purpose and 30% was kept aside as a totally unseen test set instances in mutually exclusive manner to ensure unbiased performance testing with no scope for memory from data instances.

Besides raising the trained models and testing it, as mentioned in the above section, 10 times random train-test trails have also been done to evaluate the consistency of the transformer approach on Dataset “A” and “B”. For every such trial, a total of 3,978 and 2,746 plant pre-miRNAs and equal number of negative instances, respectively, formed the training datasets. A total of 3,412 (Dataset “A”) and 2,354 (Dataset “B”) instances formed the testing datasets. In this entire study this was also ensured at every stage that no instances overlapped between train and test sets in order to avoid any chance of bias and memory. 10 folds random trials were performed where the train and tests instances from Dataset “A” and “B” were selected randomly and in mutually exclusive non-overlapping manner. Every time the model was built from the scratch using the training data and was tested upon the corresponding test set. This 10-fold random trials concurred with the above observed performance level and scored in the same range consistently. The difference between train and test mean absolute error (MAE) across 10-fold random validation trials was in the range of 0.009 to 0.0132 (Dataset “A”) and 0.007 to 0.0106 (Dataset “B”) which indicates the model was trained well with no significant overfitting. All of them achieved good quality ROC curves with high AUC values in the range of 0.9294 to 0.9436 (Dataset “A”) and 0.9734 to 0.9779 (Dataset “B”) while maintaining reasonable balance between specificity and sensitivity. (Supplementary Table S3 Sheet 4-5). As emerges from the performance metrics evaluation for the build model and their AUC/ROC plots (Supplementary Figure S2a and b), the developed transformer based pre-miRNA classification approach scored high on performance with consistent and reliable performance.

Integrating Transformer as a trainable feature extractor works better with higher dimensions and instances to learn from. In the input layer combined encodings for sequence and structure derived words were used on which the Transformer block gave remarkable results. The 14 multi-head attention system ensured proper attention to each word while mitigating any chance of wrong weighting and association mapping between the words while taking care of right context in the sentence.

### miWords consistently outperforms all the compared tools for pre-miRNA discovery

This study has performed a series of different comparative benchmarkings. The first two are covered in this section. In this comparative benchmarking, the performances of eight compared software were studied across the Datasets “A”, “B”, and “C”. The compared tools covered the classical machine learning as well as recently developed Deep Learning approaches for pre-miRNA discovery: miPlantPreMat (SVM based), HuntMi (Ensemble method of Random Forest), PlantmirP-Rice (Ensembl method of Random Forest) (57), microPred (SVM based), plantMiRP (SVM based), mirDNN (convolutional deep residual networks), deepMir (CNN based) and deepSOM (deep learning based SOM). Besides this, the benchmarking has also considered three different datasets to carry out a fully unbiased assessment of performance of these tools across different datasets. The first dataset considered was the testing dataset part of Dataset “A” and “B”. Besides measuring the performance of miWords of this neutral and totally unseen testing part of Dataset “A” and “B”, performance of the other eight tools was also benchmarked. The performance measure on the test set of Dataset “A” and “B” gave an idea how the compared algorithms in their existing form perform.

The third dataset “C” was used to carry out objective comparative benchmarking, where each of the compared software was trained as well tested across a common dataset in order to fathom exactly how their learning algorithms differed in their comparative performance.

All these eight software were tested across both the datasets which covered more than 27 plant species considered where miWords outperformed all of them across both the datasets, “A” and “B”, for all the performance metrics considered (Figure 5a and b). As already reported above for the Dataset “A” and “B” test set, miWords scored the accuracy of 94.08% and 98.04% with MCC value of 0.88816 and 0.9609, respectively, while displaying a very good balance between sensitivity and specificity with difference of just 0.4% on Dataset “A” and 1.0% on Dataset “B”. On the same Dataset “A” and “B”, the second best performing tool was plantmiRP-Rice which scored an accuracy of 87.89% and 92.56%, respectively, and MCC values of 0.75 and 0.8560, respectively, values far behind the values observed for miWords. It displayed a gap of just 0.76 (Dataset “A”) between sensitivity and specificity, a gap which is slightly higher than miWords, but yet a good balance between sensitivity and specificity. A Chi-square test confirmed that miWords significantly outperformed the second best performing tool on Dataset “A” comparative benchmarking (p-value<<0.01).

**Figure 5:**
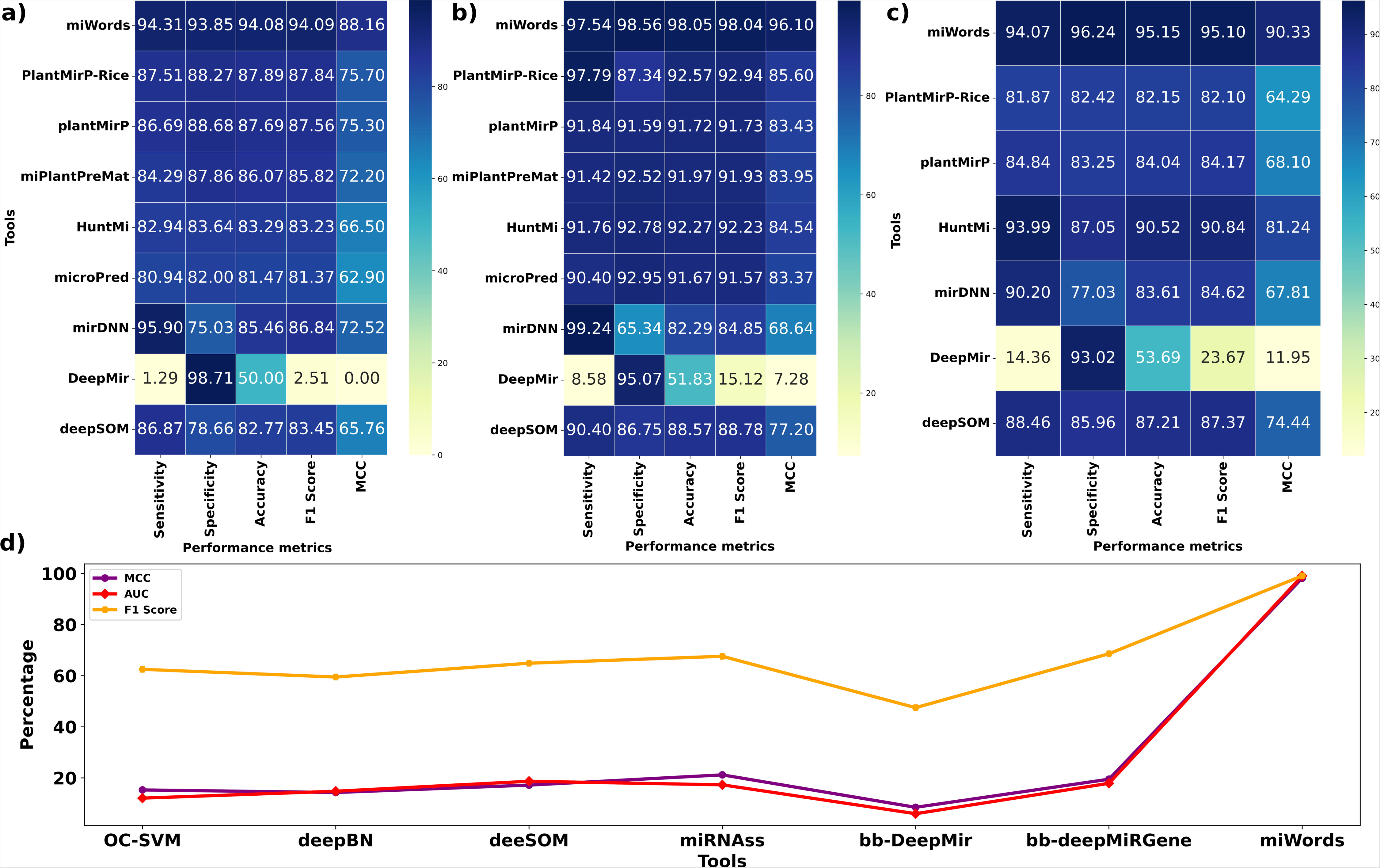
Comparative bench-marking results for miWords for three different datasets. **a)** Bechmarking result on Dataset “A”. Here all the compared tools were tested on the testing dataset part of the Dataset “A” which was totally unseen and untouched for all the compared tools including miWords. This gives a view of how the compared software would behave in their existing form and models. b) Bechmarking result on Dataset “B”. This datasets contained the high confidence refined and filtered entries from miRBase while taking support and evidences from three other databases (sRNAanno, PmiREN, and PNRD). **c)** Objective comparative benchmarking on Dataset ”C”. Here, all the compared tools were first trained on a common dataset for training and then tested on a common mutually exclusive dataset for their performance. This gave a clear view on the performance of each of the compared algorithms. (d) Comparative benchmarking done on the imbalanced dataset introduced by Bugnon et al (46). All the compared tools were trained and tested on this common data-set for objective comprative benchmarking for imbalanced dataset performance. The logic for such dataset is that in usual genomic annotation conditions, the negative instances are manifold higher than the pre-miRNA regions. A capable software should perform good on such imbalanced dataset. Here also miWords outperformed all the compared software. From the plots it is clearly visible that for all these datasets and associated benchmarkings, miWords consistently and significantly outperformed the compared tools for all the compared metrics.

On the Dataset “C”, all these tools were trained on the same common training dataset and tested across the common testing dataset in order to achieve the objective comparative benchmarking of the algorithms. However, two tools, microPred and miPlantPreMat could not be included in this part of bench-marking as both these tools don’t give provision to train on another dataset and rebuild models. Thus, in this part of benchmarking, the remaining six tools and miWords were trained and tested on the common dataset. In this benchmarking also miWords outperformed all the compared tools with significant margin with the similar level of performance (Figure 5c). miWords clocked an accuracy of 93.6% and MCC of 0.87, while displaying a good balance between sensitivity and specificity where gap of only 0.94% was observed. The second best performing tool was HuntMi which attained an accuracy of 90.5% and MCC value of 0.81 but displayed much higher gap of ∼7% between sensitivity and specificity scores, exhibiting significant performance imbalance. A Chi-square test done here also confirmed that miWords significantly outperformed the second best performing tool, HuntMi (p-value<<0.01).

Besides this all, one more interesting objective comparative benchmarking analysis was done on an imbalanced dataset recently provided by Bungan et al, 2021 (42). In their benchmarking study, they knowingly created an imbalanced dataset with much higher negative instances to mimic the actual genome condition where class imbalance is pronounced. There they strongly attracted the attention on the fact that how most of the existing pre-miRNA discovery software performed very poorely.

This dataset contains 839 pre-miRNA (positive) and 13,56,616 (negative), utilizing this dataset an imbalanced dataset was created in the ratio of 1:1616. The dataset was split into 70:30 ratio to train and test the model for miWords, maintaing 1:1616 ratio of positive and negative instances. miWords’s performance was compared fot this dataset also where the compared tools were also trained and tested on this common dataset. Here too miWords scored the highest for all the performance metrics with big lead margin than the rest of the compared six software. miWords achieved an MCC of 98.18%. (Figure 5d).

Also this needs to be noted in both the bench-marking tests, miWords scored much higher MCC values which suggests consistent and robust performance. MCC gives high score only when a software scores high on all the four performance parameters (true positive, false positive, true negative, false negative). As it is visible from the score distribution for all the metrics (Figure 5), miWords also exhibited least dispersion among all. miWords’s performance points out that more appropriate features may be learned through training on syntax of words, and their subsequent efficient encoding with multi-headed attention using Transformer as an encoder. The full details and data for this benchmarking study are given in Supplementary Table S4 Sheet 1-4.

### Genomic context learning on transformer scores delivers extremely good results on genome wide annotaitons of pre-miRNAs

Performance over standard testing datasets may be claimed good, as has been done by most of the published software in the past. But in actual application of genome annotation, huge performance gaps exist and far below the acceptable limits. Some recent reports have highlighted that how much poor performing most of the existing pri-miRNA discovery tools become in the real situation applications like genomic annotations where most of them end up reporting very high proportion of false positives (3, 42). This has also led to one of the rare event of mass withdrawal of entries of plant miRNAs from databases like miRBase, recently. Taking note of such extreme events in plant miRNA biology, a very insightful commentary was made (4). There they recommended some protocols to identify genuine pre/miRNAs candidates and suggested a necessary run against some well studied established genome like *Arabidopsis* to compare how much false positive identifications were made by any plant miRNA discovery tool. It has become the standard protocol to assess the success of such tools in real application of locating pre-miRNA regions across a genome. Most of the existing tools perform highly unreliably in genome annotation. The best performing tools were found to be necessarily dependent on sRNA sequencing read data to identify pre/miRNAs and yet they end up with lots of false identifications as transpires in discussion below also.

One important problematic factor about all these existing tools is that they hardly acknowledge the role of relative information from the flanking non-miRNA regions in accurate identification of miRNA regions during genome scanning. Their core pre-miRNA discovery algorithms hardly consider such factors. While in actual the relative scoring patterns between miRNA regions and neighbourhood non-miRNA regions can become a highly informative for more accurate discrimination. We hardly found any study which performed genomic scanning and tried to further learn on the scoring patterns for pre-miRNA regions and non pre-miRNA regions. A high scoring pre-miRNA region is expected to display higher scoring distribution across its bases along with gradual decline when compared to its non-miRNA flanking regions where scoring is also expected to exhibit random and sharper trend. Doing so would also help in detecting boundaries of the pre-miRNA regions. A t-test between the flanking regions T-score distribution and pre-miRNA regions supported this view (p-value < 0.05). Thus, it became another important aspect of miRNA regions and to refine their discovery in genomic context. Therefore, for the first time we conducted such study and trained another deep learning CNN based module on the obtained T-scoring profiles in actual run across the genomes.

For T-scoring based CNN two different models were raised from the Dataset “A” (called Model “A” for T-Score CNN) and “B” (Called Model “B” for T-Score CNN). In model “A”, a total of 604 *Oryza sativa* pre-miRNAs regions were considered for raising T-scoring CNN model. In model “B”, a total of 631 pre-miRNAs regions were considered from species *Arabidopsis thaliana, Oryza sativa, Zea mays, and Glycine max* for raising T-scoring based CNN model.

In this process, the transformers, which were trained and raised in the previous step, were run across the annotated genome of *Arabidopsis thaliana, Oryza Sativa, Zea mays, and Glycine max* where the transformer’s scoring patterns was recorded for the annotated pre-miRNAs regions and the non-pre-miRNA regions. This was done with a sliding window of size 200 bases, which generated vectors of transformer scores per base for the pre-miRNA regions and non-miRNA regions. The obtained scoring profiles for pre-miRNA regions constituted the positive datasets while the T-score distribution per base for the flanking 500 bases both sides of pre-miRNA regions having no pre-miRNAs constituted the negative dataset. In the methods section above, full details of implementation for this module is already given. The position specific scoring profiles were converted into a matrix of 280×10 dimension, where the rows contained the scoring values in the range of 0-1 in a discrete manner, while the columns captured the base position for the given window size. One-hot encoding was done in this matrix where for any given base position, the corresponding value was assigned value from the range of 0-1. This mimicked a pixeled image which now could be passed through Convolution neural nets (CNN). CNN has been brilliant in recognizing spatial patterns, and we expected them to capture the assumption that pre-miRNA regions display a significantly different scoring pattern for the bases than those belonging to non-miRNA regions. This way for model A, a total of 604 experimentally validated *Oryza sativa* pre-miRNA instances belonged to the positive instances dataset, and a total of 604 non-miRNA regions (comprised of randomly selected 5’ and 3’ flanking non-miRNA neighbors) belonged to the negative dataset. This dataset was split into a ratio of 70:30 to train and test the CNN. Likewise, for Dataset “B” derived model, a total of 631 pre-miRNAs from *Arabidopsis thaliana*, *Oryza sativa, Zea mays, and Glycine max* belonged to the positive instances dataset, and an equal number of non-miRNA regions (comprised of randomly selected 5’ and 3’ flanking non-miRNA neighbors) belonged to the negative dataset. This data too was split in the ratio of 70:30 on which the CNN was trained and tested. For Model “A”, an accuracy of 78.6% with sensitivity 79.21% and specificity 77.99% was observed. When the same test was carried out for the Model “B”, which had more refined high confidence data-set, the accuracy of T-Score CNN attained 90.76% with 89% sensitivity and 92.73% specificity. The clear benefit of having the high-confidence instances in Dataset “B” reflected here too. Also, from here it transpires that if the Transformer-XGBoost classification system’s scoring scheme is learned with genomic context using the above mentioned T-Score CNN, the actual application in genomic annotations would benefit a lot.

Therefore, now the real test was to check the raised model’s performance was on its ability to correctly identify the miRNAs and non-miRNA regions in some very well annotated and studied genome. This would provide the clear picture about the performance of such software in their practical application of discovery of miRNA regions across genomes. For this, the *Arabidospsis* genome was taken with its full annotations. *Arabidopsis* has genome size of 119.763 MB and a total of 326 pre-miRNAs are reported for *Arabdiopsis thaliana* in miRBase version 22 (37). Also, this needs to be noted that in order to attain an unbiased assessment of the raised model on Arabidopsis genome, all the above mentioned model generation steps of Transformers-XGBoost and T-Score CNN were redone from the scratch with Arabidopsis data completely removed from Dataset “A” and “B” entries. The performances being discussed below for Arabidopsis genome, therefore, are from the learning which never witnessed any data from Arabidopsis previously. In the first phase, for the entire genome for each base the transformer score (T-score) was generated from two different models which were derived from the datasets “A” and “B”. This became the input to the T-scoring based CNN.

A total of 323 and 322 out of the annotated 326 pre-miRNAs of *Arabidopsis* were detected successfully from the build model derived from Dataset “A” and “B”, respectively. Hence, it was clear that discovering miRNA regions by the above mentioned system was highly accurate. The next and bigger important question was that how much novel miRNAs were identified across this genome, which could be most probably the false positive cases? A total of 771 and 759 pre-miRNAs regions were suggested by the transformers, which meant that a total of 448 and 437 false positives were called pre-miRNA regions by the transformers derived from Dataset “A” and “B”, respectively. Though this number is very much lesser than what the currently existing pre-miRNA discovery tools and approaches report (including some NGS sRNA-seq data dependent tools) (26), but even this number may be considered substantially high. However, we got an exceedingly good and surprising result when the transformers scores were passed through the above mentioned T-scoring based CNN. Just 29 (model “A”) and 43 (model “B”) novel candidates, the potentially false positive cases, were obtained for the entire *Arabidopsis* genome. This was an exceptionally good result, especially when considering the fact that it was not given even any sRNA-seq data guidance.

The existing tools which were run across *Arabidsopsis* genome with sRNA-seq read data supports identified at least 11 false positive pre-miRNA candidates and went predicting up to 12,306 miRNAs despite having sRNA-seq reads support. This clearly underlines that despite of having sRNA-seq reads data as their guide, due to inefficiency in their existing core algorithms to identify pre-miRNA regions, in actually these tools don’t benefit significantly from sRNA-seq reads data guidance and end up identifying large number of false positive instances. Also, in general they grossly missed to identify a big number of actual miRNAs with their sensitivity value ranging from 3% to utmost 86%. Figure 6 provides one such comparative benchmarking map between some of these sRNA-read data guided software performance on *Arabidopsis* genome and miWords’s relative standing with its different forms. Even without using sRNA sequencing read data unlike these category of software, miWords was found outperforming them by big margin. Thus, the implications of our developed software, miWords, are going to be high. It can identify pre-miRNA regions across the genomes highly accurately even without getting any help from sRNA sequencing experiments, directly cutting the cost of experiments, time, and efforts. miWords’s performance with sRNA-seq reads data support improved further, as transpires from the discussion below.

**Figure 6:**
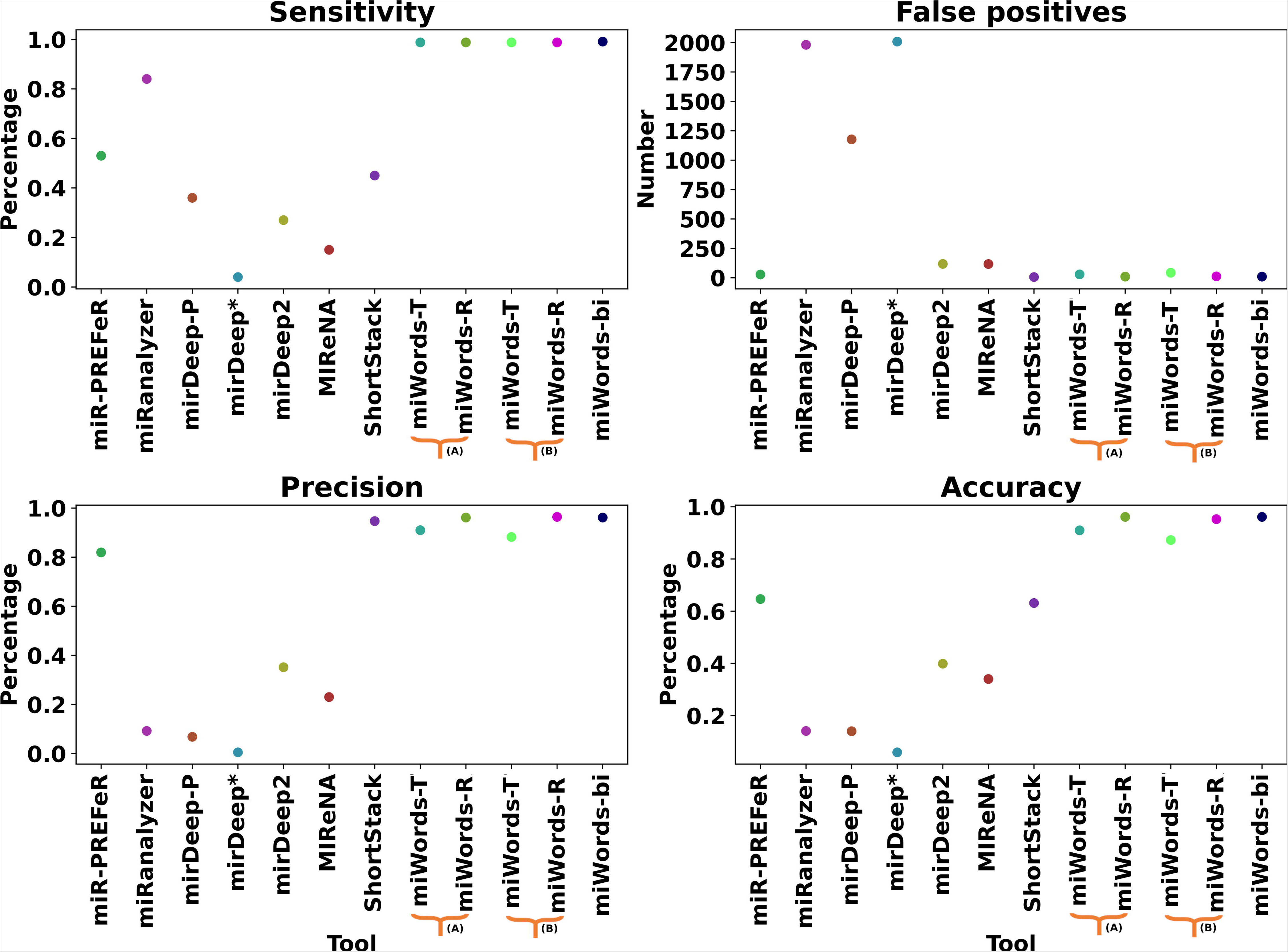
Comparative benchmarking for genomic annotation capability and performance. Most of the existing software perform poorely in the actual application of genome annotation and end up reporting large number of false positive cases. It has been recommended to assess performance of any such tool across well annotated genomes like Arabidopsis. Any reporting of novel miRNAs on such genome should be considered as a false positive case and accordingly the performance of a software may be rated. In this performance bench-marking, miWords was compared to the tools which are presently most preferred ones for genomic annotation at the present, as they use sRNA sequencing reads data as help guide to reduce their false positive predictions. As can be seen from this bench-marking plot, all the three forms of miWords (miWords-R: Working with serially connected sRNA-seq RPM CNN; miWords-T: Working without any sRNA-seq reads help and directly with genomic T-Score CNN; miWords-Bi: The bi-modal CNN form where T-score and sRNA-seq derived per base RPM representations of sequences is jointly learned outperformed all the compared tools for all these performance metrics. “A” and “B” are for the data-sets used in the present study to derive the models. In *Arabidopsis*, miWords identified all of its pre-miRNAs correctly except three of them, and reported only 10 false positives, the lowest of all.

And then naturally comes the next question: How good it would perform if some one provides sRNA sequencing data too? To answer this, miWords has implemented two different modules: 1) A serially connected CNN module, and 2) a bi-modal CNN. Both of them take Reads per million (RPM) normalized value representation per base of the sequences. As done with T-Score CNN, here also the pre-miRNA regions form the positive instances and their flanking non-miRNA regions work as the negative instances. Thus, same datasets were used here also which were used to train the T-Score CNN. T-Score CNN represented them in the form of transformer score per base, while the RPM-profile CNN here represented them in the form of normalized base coverage through sRNA-seq read data. The first one works as an additional filter to improve the result of T-Score CNN while the second one collaborates with T-Score CNN and parallelly contributes in taking the collective decision. Both of them have been trained on the above mentioned Datasets “A” and “B”. First model (model “A”) was raised on a total of 604 (433+171) positive and 604 (433+171) negative instances from *Oryza sativa* where 70% of the dataset was used to train it and 30% was used to test it. This model achieved an accuracy of 76.64% with sensitivity 77.29% and specificity 75.99%. The second model (model “B”) was raised on a total of 631 (442+189) positive and 631 (442+189) negative instance from *Arabidopsis thaliana, Oryza sativa, Zea mays, and Glycine max.* The model achieved an accuracy of 86.44% with sensitivity 84.75% and specificity 88.14%. Effect of high-confidence datasets was visible here also. Like in this entire study, here also it was ensured that no instances overlapped between training and testing to avoid any memory and bias in the learning. Details of this module is already provided in the methods section above and Supplementary Figure S3b.

The raised RPM based CNN modules were tested across the Arabidopsis genome sequences to measure how much they contribute to improve the results. As done in the previous steps of T-Score CNN, here also the Arabidopsis sequences were completely removed from the training and testing datasets to ensure totally unbiased measure of the performance without any scope of memorization of instances and data from Arabidopsis. On passing through this optional and serially connected RPM-CNN module the false positive identification further decreased to just 10 cases (model “A”) and 12 (model “B”), a number much lesser than the number of 28 false positive cases reported by best performing software, miR-PREFeR after taking support from sRNA-seq data. Figure 6 provides the comparative benchmarking of miWords and its various versions with seven best performing tools which use sRNA-seq data, clearly suggesting the top notch performance by miWords.

This also may be noted that in a previous bench-marking study on these compared software, it was found that they are sensitive towards the size of sRNA-seq data and number of studies included. As this data volume and number of studies increases, the number of potential false positives by these software was reported to increase also (25). The reported total number of total novel 28 miRNAs by the best performing tool, miR-PREFeR, was when only two experimental conditions sRNA-seq data were considered. As soon as they considered higher number of samples (6), their reported novel miRNAs number shot up to 49. The same trend was observed for almost all of the compared tools with much higher off-shooting. In the present study we had considered comparatively much bigger sRNA-seq data for *Arabidopsis*, a total of 88 samples, and yet did not see such overshooting effect and reported only 10 and 12 novel pre-miRNAs for the RPM based CNN module derived from Dataset “A” and “B”, respectively, as mentioned above. Even without sRNA-seq data support and just based on its Transformer scoring CNN module, miWords outperformed all of those compared software which needed sRNA-seq data essentially. As already shown above, the core transformer based part of miWords, outperformed all the compared software with large margins, which only further improved with addition of T-Score CNN and sRNA-seq data guided RPM-CNN which incorporated relative genomic context information and transcriptional information.

The above mentioned sRNA-seq RPM based module was implemented serially where output from the T-score CNN is passed on to further filter and refine the result in optional manner, depending upon the availability of sRNA-seq data. Additionally, a bi-modal CNN was also implemented which takes the inputs in the two forms parallelly: genomic T-scoring values and sRNA-seq derived RPM sequence representation of the corresponding sequence. The same training and testing datasets were used as mentioned above, while keeping the same splitting approach. It differed from the above mentioned sequential approach that here it is not an optional approach and it works along with the transformer scoring sequences to directly classify the pre-miRNA regions in genomic context, instead of being an additional refinement/filtering step. The model achieves an accuracy of 88.54% with sensitivity 87.67% and specificity 89.41% while training and testing on the above mentioned Dataset “B”, performance 2% higher than the serial mode. Like done before, here also this model was retrained on the dataset where all instances from *Arabidopsis* were removed, in order to ensure unbiased performance without any scope of memorization of *Arabidopsis* instances. This parallel bi-modal CNN performed slightly better than model “B” serial mode while detecting 323 out of 326 pre-miRNAs and just 10 novel pre-miRNAs across the Arabidopsis genome, showcasing the combined parallel learning on T-scores sequences with sRNA-seq derived RPM sequences delivers better performance than the serial filtering arrangement as described above. However, to run this bi-modal CNN, one would essentially require sRNA-seq data and generate the RPM representation for each base, while the previous one gives flexibility to work even without the sRNA-seq read data, while achieving far superior result than the existing pool of those software which essentially require sRNA-seq data to identify miRNA regions.

Thus, miWords emerged not only as the best performing software, but has exceptionally exceeding performance with much stability and reliability. It has emerged as the most suitable software to annotate plant genomes for miRNA encoding regions.

### Application: Using miWords to annotate pre-miRNA regions across Tea genome

To exhibit the applicability of miWords in practical scenario of genome scanning for pre-miRNA discovery, miWords was run across the *C. sinensis* genome whose size is 3.06 GB. Tea is the most consumed beverage, a highly important commercial crop with medicinal values also, and which is also highly sensitive to climate change. Though its genome has been revealed, to this date there is no entries for Tea miRNAs in miRBase. Thus, reporting miRNAs of Tea here would not only exhibit the application demonstration of miWords in genome annotations for miRNAs, but also benefit the research groups working on Tea to understand its molecular systems.

The first run of miWords, which was the transformer part (Dataset “A” and “B”), identified 17,044 and 16,676 pre-miRNA regions in the tea genome while basing upon Dataset “A” and “B”, respectively. For model “A” (derived from Dataset “A”), scoring for all these regions were passed to the T-scoring profile based CNN module, which screened them and reported a total of 3,194 pre-miRNA candidates. Finally, the sRNA-seq data supported RPM-CNN module was run on it, which reported a total of 821 pre-miRNA candidates. For this part of the run, we had collected sRNA-seq reads from 104 samples while covering 34 different conditions. This is not necessary that all other discarded pre-miRNA candidates reported by the previous step were false positive, as sRNA-seq data are highly condition specific, and many conditions may not have been captured by the existing sRNA-seq experiments done on Tea. Yet, these 821 pre-miRNAs from Tea genome may be considered as the most confident cases.

Likewise, 16,676 pre-miRNA regions in the tea genome which were detected from transformers (Dataset “B”) were scanned through the bi-modal CNN system too. bi-modal CNN reported a bit lesser number of 803 pre-miRNAs candidates in Tea (Figure 7). Between the serially arranged RPM-CNN’s identified 821 pre-miRNAs and 803 pre-miRNAs detected through the bi-modal CNN, 788 regions were common. All of these potential pre-miRNA regions exhibited sRNA-seq reads data mapping to them across multiple samples where at least five reads mapped in each condition. 10 of these pre-miRNA regions were also re-validated using qRT-PCR for their being transcriptional status where almost all of them were found expressing. Figure 8 shows their qRT-PCR expression levels, their secondary structures, and the highlighted box regions for which dominant sRNA-seq reads data were found mapping. Complete Tea miRNAome details are given in the Supplementary Table S4 Sheet 5. All this gave strong support evidence to the identified pre-miRNAs.

**Figure 7:**
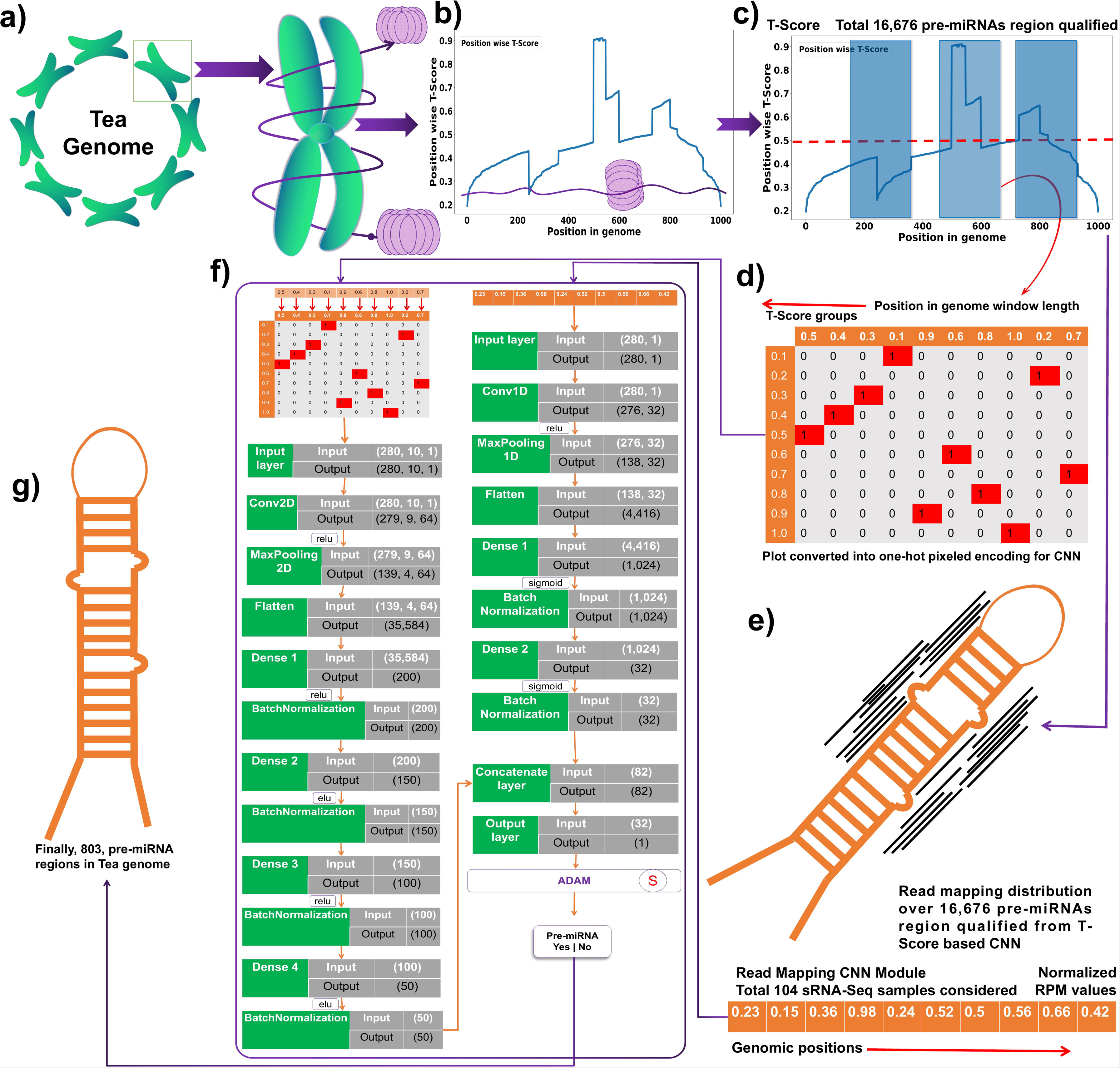
Details of the workflow carried out to annotate the Tea genome for pre-miRNAs. A total of 803 pre-miRNAs were identified in *C. sinensis* using the bi-modal CNN form which used the Transformer scoring for each base and sRNA-seq reads mapping information in the form of RPM per base. All of the identified miRNA regions had sRNA-seq reads supports from multiple experimental conditions.

**Figure 8:**
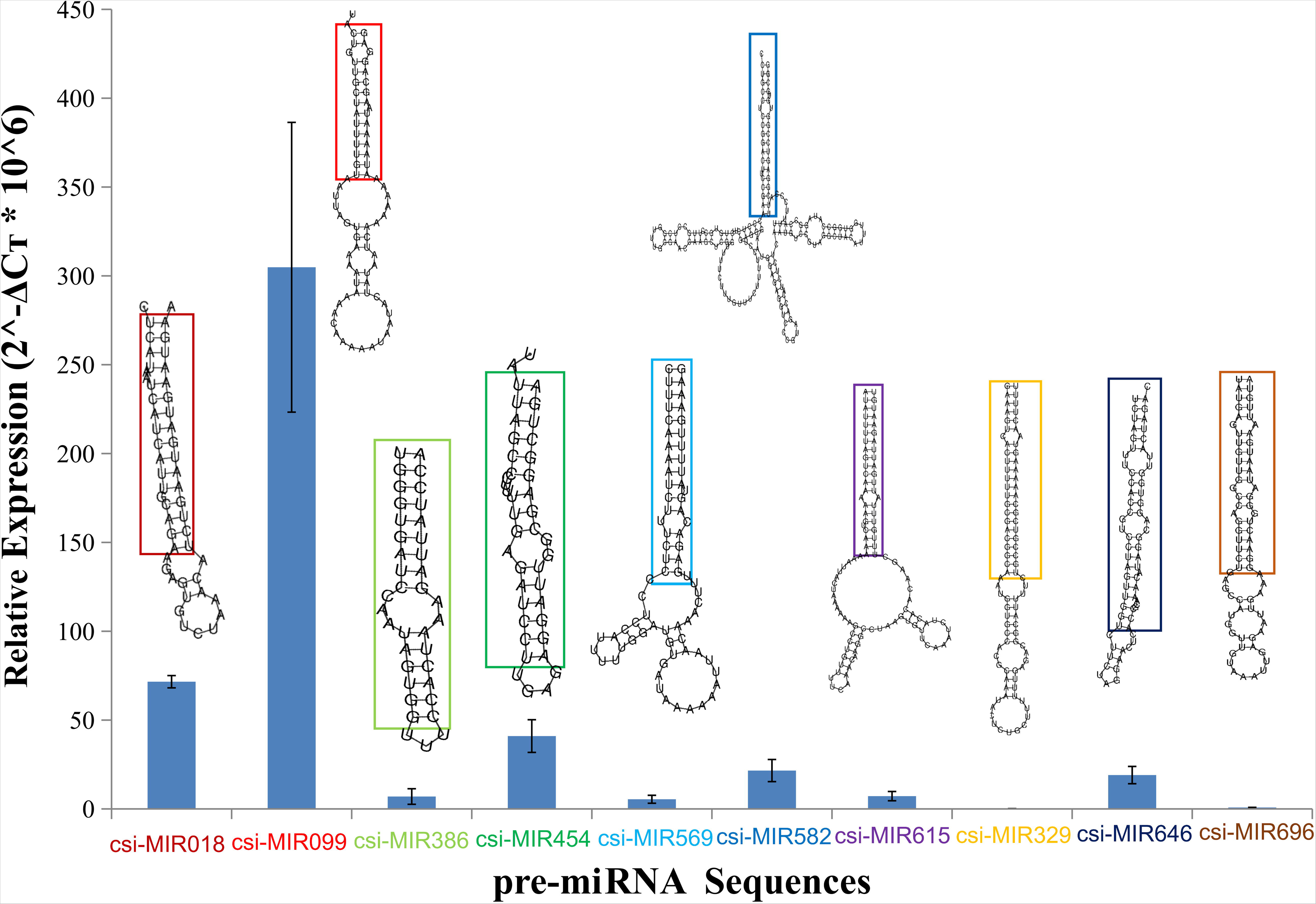
Experimental re-validation of the identified pre-miRNA regions in *C. sinensis* for their transcriptional activity. miRNA expression analysis of selected 10 pre-miRNAs by quantitative real time-PCR in tea leaves (*Camellia sinensis*). pre-miRNAs expression is presented relative to 18S rRNA. Mean ± SD of triplicate quantitative real-time PCR from a single cDNA sample. The upper part shows the corresponding secondary structure for the sequence, while the colored box shows the region for which sRNA-seq reads were found mapping dominantly.

### Webserver and standalone implementation

miWords has been made freely available at https://scbb.ihbt.res.in/miWords/ as a very simple to use webserver. The server has been implemented using Plotly visualization library, Python, Javascript, PHP, Shell, and HTML5. The user needs to paste the RNA/DNA sequences in FASTA format into the text box or upload the RNA/DNA sequences in FASTA format and then click the submit button for the identification. After a while the result page appears from where results can be downloaded in a tabular format.

However, in actual situation like whole genome sequence scanning, web servers are practically not suitable and one needs standalone version mainly. For heavy duty real scenario application like genome annotation, a standalone version of miWords has also be provided via a link tab on the same server page or can be downloaded from github. The provision to see the results in plots is also given there.

## Conclusion

miRNAs define one of the largest regulatory systems of eukaryotes. The mature miRNAs are processed out of their precursors (pre-miRNAs). Though lots of software have been developed to identify pre-miRNAs as their discovery is the core of miRNA biology, in actual practical application of discovery of pre-miRNAs across genomes these software remain far away from the acceptable limits. This is much more grave when dealing with plant genomes where the generally considered properties and features to distinguish a pre-miRNA regions don’t work as a strong discriminator. The present work has approached pre-miRNA regions as a sentence hidden within genome where the relative arrangements of the words in the form of dinucleotides, trimers, pentamers, and structural triplets define a sentence. Once the syntax is there, one also needs they are read by an intelligent reader which could decode the relationship within the words. This was achieved by a revolutionary deep-learning algorithm of transformers which assigned contextual attention scores to the words withing the sentence using 14 multi-headed transformers which finally passed their learning to an XGBoost classifier to generate classification scores. The next part of this system, miWords, applied a convolution networks system which learned from the transformer-scoring pattern across the genome and partitioned it into the pre-miRNA and non-pre-miRNA vicinity regions to successfully define the boundaries and make it possible to run such software for its practical utilities for genomic annotation. Total four direct and different benchmarking studies were carried out in this study involving more than 10 different published software to identify miRNA regions, and in all of them miWords significantly outperformed the compared tools. This also included the class of software which are prime choice for genomic annotation for miRNAs, the Next-gen sequencing reads data guided software. Even without using its sRNA-seq guided module, miWords outperformed software which essentially need sRNA-seq guidance, making it the most suitable and accessible software for genome annotation for miRNAs which can work with much higher accuracy than others even without cost and time on running NGS experiments. Additionally, miWords, also provides an optional module to use sRNA sequencing reads data to further refine the results. As an application demonstration, miWords was run across the Tea genome no identify pre-miRNAs across its genome where it reported 803 pre-miRNA regions, all supported by sRNA-seq reads for multiple conditions as well as found transcriptionally active through qRT-PCR experiemnts also. This all has validated the approach of miWords and its capabilities to annotate genomes for miRNAs. miWords appears to be the most capable tool which can solve the long pending quest for a software which could be reliably used for genomic annotation for miRNAs.

## Declarations

### Availability of data and materials

All the secondary data used in present study were publicly available and their due references and sources have been provided in the Supplementary Table S1-4. The software has also been made available at Github at https://github.com/SCBB-LAB/miWords as well as at https://scbb.ihbt.res.in/miWords/.

### Authors’ contributions

SG carried out the computational part and benchmarking of the study. RS conceptualized, designed, analyzed and supervised the entire study. VS and RK carried out the wet lab experiment. RK supervised the wet lab experiment. SG and RS wrote the MS.

### Ethics approval and consent to participate

Not applicable.

### Consent for publication

Not applicable.

## Supplementary information

**Supplementary Figure S1: Optimization results for hyperparameters for transformers part of the hybrid Transformer-XGBoost model. A)** Batch size optimization, **B)** Dropout rate optimization, **C)** Second Dropout rate optimization, **D)** Learning rate, **E)** Number of units per dense layer 1, **F)** Epoch size optimization **G)** Embedding size, **H)** Number of units per dense layer inside Transformer, **I)** Number of dense layers 2, and **J)** Number of Attention heads.

**Supplementary Figure S2: AUC/ROC plot for Ten fold cross validation.** The AUC/ROC plots for the hybrid models for the testset clearly showcase the robustness and highly reliable performance of the implemented hybrid Transformer-XGBoost model. **a)** AUC/ROC plot for the hybrid models for the Dataset “A” testset and **b)** AUC/ROC plot for the hybrid models for the Dataset “B” testset.

**Supplementary Figure S3: Detailed pipeline of the scoring and RPM profile. A)** The image provides the outline of CNN architecture implemented for scoring based profiles for better classification of pre-miRNAs. **B)** Architecture of RPM based CNN model implemented second level classification of pre-miRNAs.

**Supplementary Table S1: Stats of different properties of pre-miRNAs between plant and animal species.**

**Supplementary Table S2: sRNA-seq data description for *Arabidopsis thaliana* and *Camellia sinesis* and species wise breakup for Dataset A.**

**Supplementary Table S3: Evaluation and optimization of hybrid Transformers-XGBoost model.**

**Supplementary Table S4: Comparative benchmarking results of miWords.**

## Acknowledgments

The work was carried out under the aegis of The Himalayan Centre for High-throughput Computational Biology (HiCHiCoB), a BIC supported by DBT, Govt. Of India. We are thankful to CSIR for the funding support they gave for this project. SG is thankful to CSIR and DBT, India for financial support as project associateship. SG and VS are thankful to Academy of Scientific and Innovative Research (AcSIR) for their PhD enrollment. This MS has CSIR-IHBT MSID #5137.

