## Supplementary figures and images for "Sentences, Words, Attention: A “Transforming” Aphorism for the Discovery of pre-miRNA Regions across Plant Genomes"

### Supplementary Figure S1

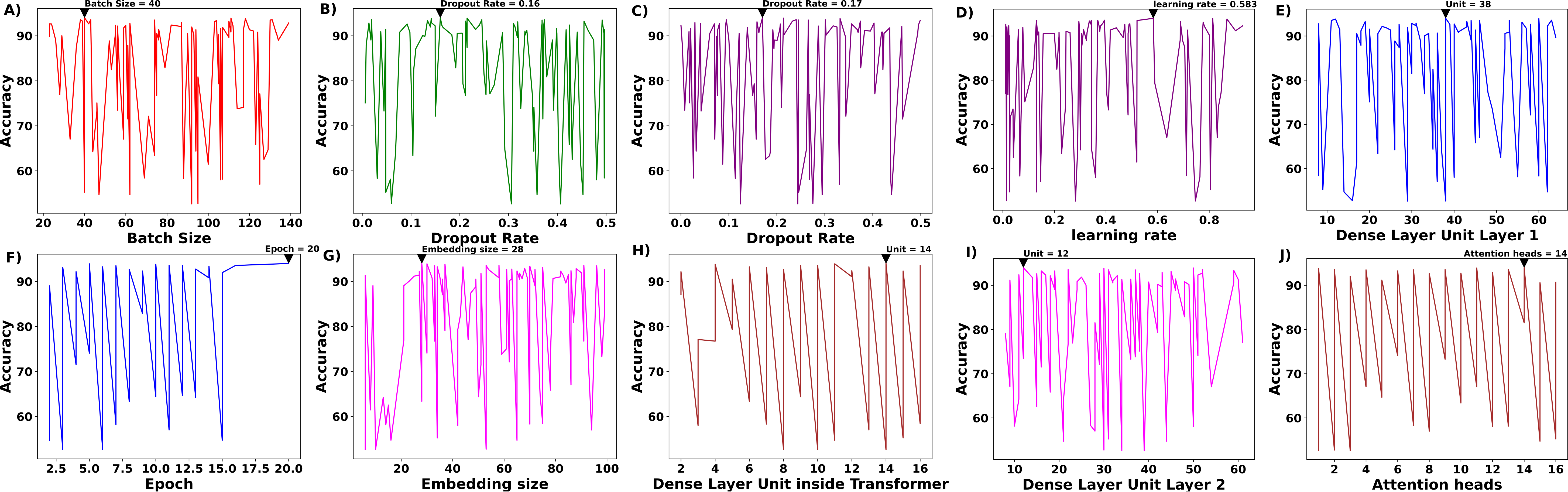

### Supplementary Figure S2

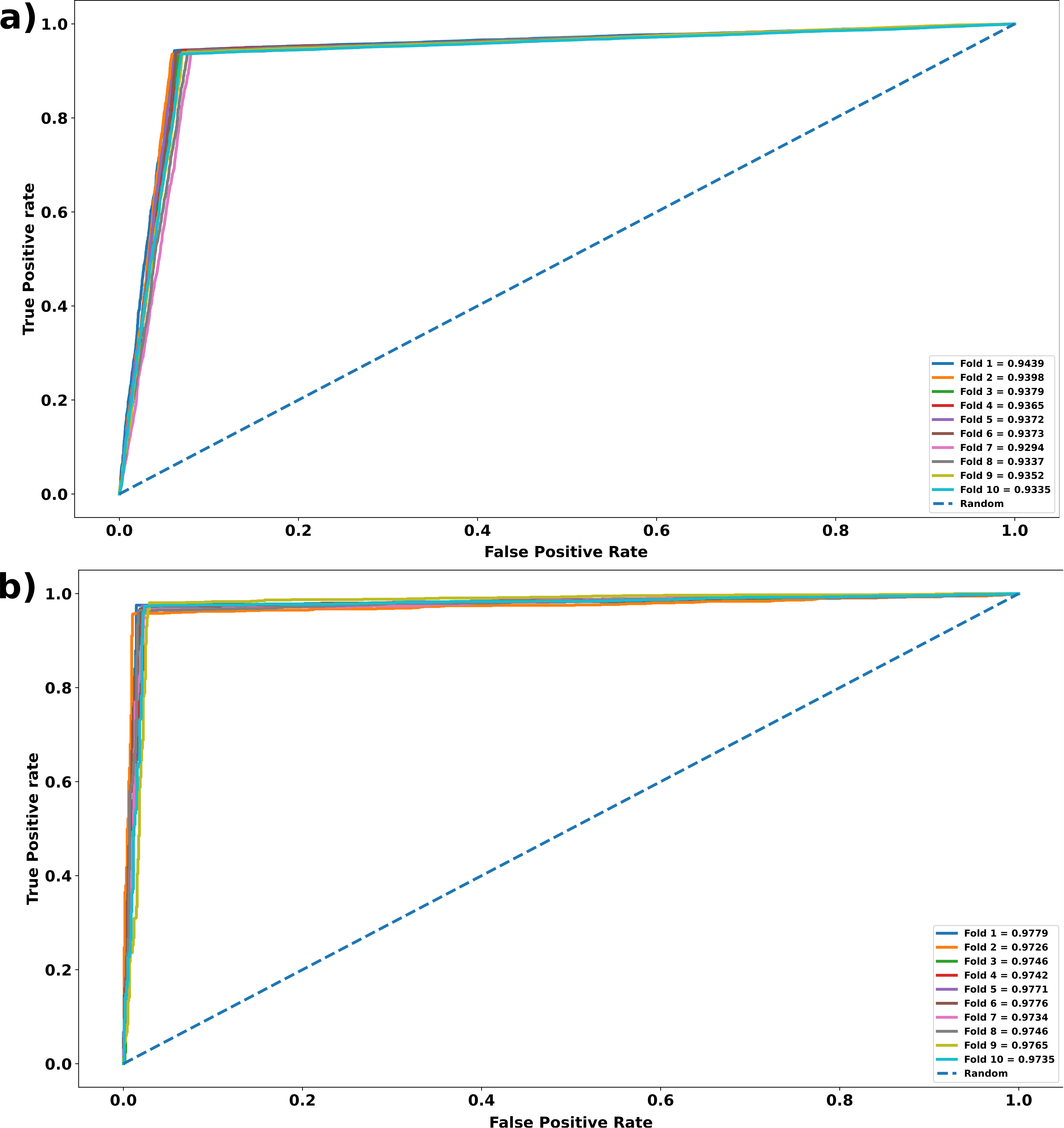

### Supplementary Figure S3

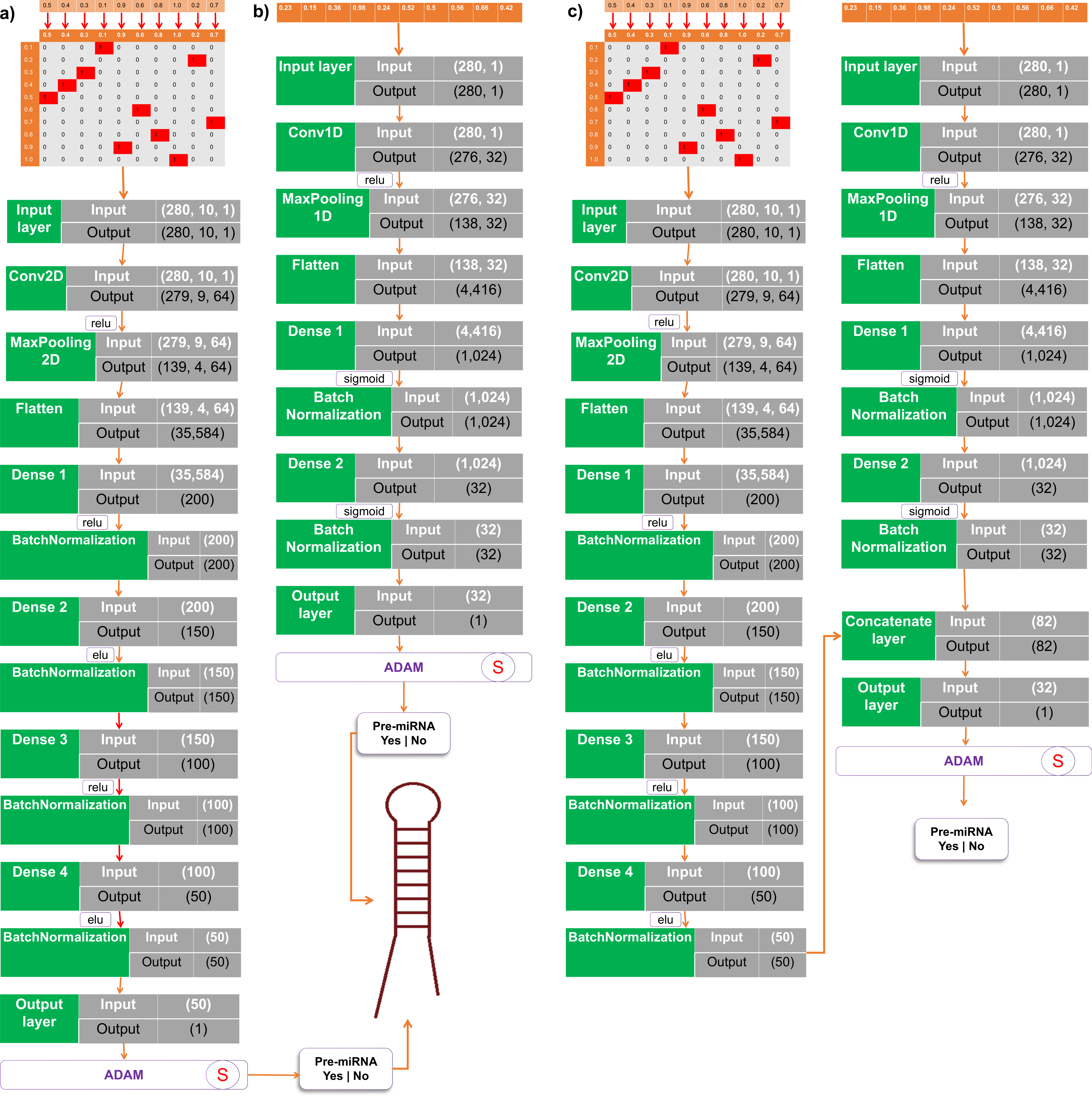
